# Evolution of a large periplasmic disk in *Campylobacterota* flagella facilitated efficient motility alongside autoagglutination

**DOI:** 10.1101/2023.09.08.556628

**Authors:** Eli J. Cohen, Tina Drobnič, Deborah A. Ribardo, Aoba Yoshioka, Trishant Umrekar, Xuefei Guo, Jose-Jesus Fernandez, Emma Brock, Laurence Wilson, Daisuke Nakane, David R. Hendrixson, Morgan Beeby

## Abstract

Although the bacterial flagella of *Escherichia coli* and *Salmonella enterica* are distributed around the cell body, many bacteria instead place their flagella at their poles. This widespread form of flagellar motility is relatively poorly understood, but these polar flagellar motors invariably feature periplasmic disk structures of unknown function. The flagellar motor of *Campylobacter jejuni* features a 100 nm-wide periplasmic disk associated with scaffolding a wider ring of motor proteins to increase torque, but the size of this disk is excessive for a role solely in scaffolding motor proteins. Here we show that the basal disk in *C. jejuni* is a flange that braces the motor during disentanglement of the flagellar filament from interactions with the cell body and other filaments, interactions that are otherwise important for host colonization. Our results reveal an entanglement of co-dependencies in the evolution of flagellar motor structure and cell plan in the Campylobacterota (previously epsilonproteobacteria). Note that this manuscript has a sibling manuscript titled *’Molecular model of a bacterial flagellar motor in situ reveals a “parts-list” of protein adaptations to increase torque’* that describes a molecular model of the *Campylobacter jejuni* flagellar motor discussed here.

## Introduction

Many Bacteria use flagella, membrane-embedded rotary motors connected to external helical propellers, to move through their environments^1^. All flagella, across genera separated by hundreds of millions of years of evolution, share the same core proteins. A ring of motor proteins called stator complexes harness ion flux to rotate a tens-of-nanometres-wide cytoplasmic ring, or C-ring. The C-ring is connected to a chassis structure in the inner membrane called the MS-ring which, in turn, forms the base of the ∼30-nm axial driveshaft which spans the periplasm (the rod), the ∼50-nm universal joint for torque redirection (the hook), and the multimicron-long flagellum itself. Transiently reversing motor rotation causes the cell to randomly reorient; chemotaxis is achieved by inhibiting tumbling if the chemical environment is becoming more favourable.

Research has focused on the model organisms *Escherichia coli* and *Salmonella typhimurium*, which have several flagella distributed across the side of their rod-shaped cells^2^. When all flagella rotate counterclockwise (CCW), they form a coherent bundle that propels the cell. The low-complexity architecture and randomized placement of flagella in these organisms, however, is only one paradigm of flagellation. Many species, including from *Vibrio*, *Pseudomonas*, *Bdellovibrio*, *Helicobacter*, and *Campylobacter* genera, assemble motors at one or both poles. Polarly-localized flagellar motors are more structurally complex than lateral flagella, but the significance of their increased complexity is unclear.

Polar motors preclude filament bundling as a swimming style. Swimming in bipolar- flagellated species involves wrapping the leading flagellum around the cell body to exert thrust in in concert with the lagging flagellum^3–6^. *Campylobacter jejuni*, a member of the *Campylobacterota* (previously epsilonproteobacteria^7^), reorients by pulling its wrapped (leading) filament from the cell surface by transiently switching motor rotation to clockwise (CW); this allows the previously unwrapped (lagging) filament to take its place wrapped around the cell body from the other pole.

*C. jejuni* has one of the largest and most complex flagellar motors, including several periplasmic disk structures not seen in *Salmonella*^8^ (Fig. 1A). The largest is the basal disk, which assembles from thousands of copies of the lipoprotein FlgP to over 100 nm wide. The basal disk is required for assembly of other periplasmic disks that form a scaffold required for incorporation of stator complexes into the motor. This periplasmic scaffold facilitates positioning of a wider ring of additional stator complexes, consistent with the *C. jejuni* motor producing approximately three times the torque of the *Salmonella* motor^8^. The basal disk, however, is wider than the periplasmic scaffold and stator complex ring, and it is unclear what benefit *C. jejuni* gains from such an apparently excessively large disk.

**Figure 1:**
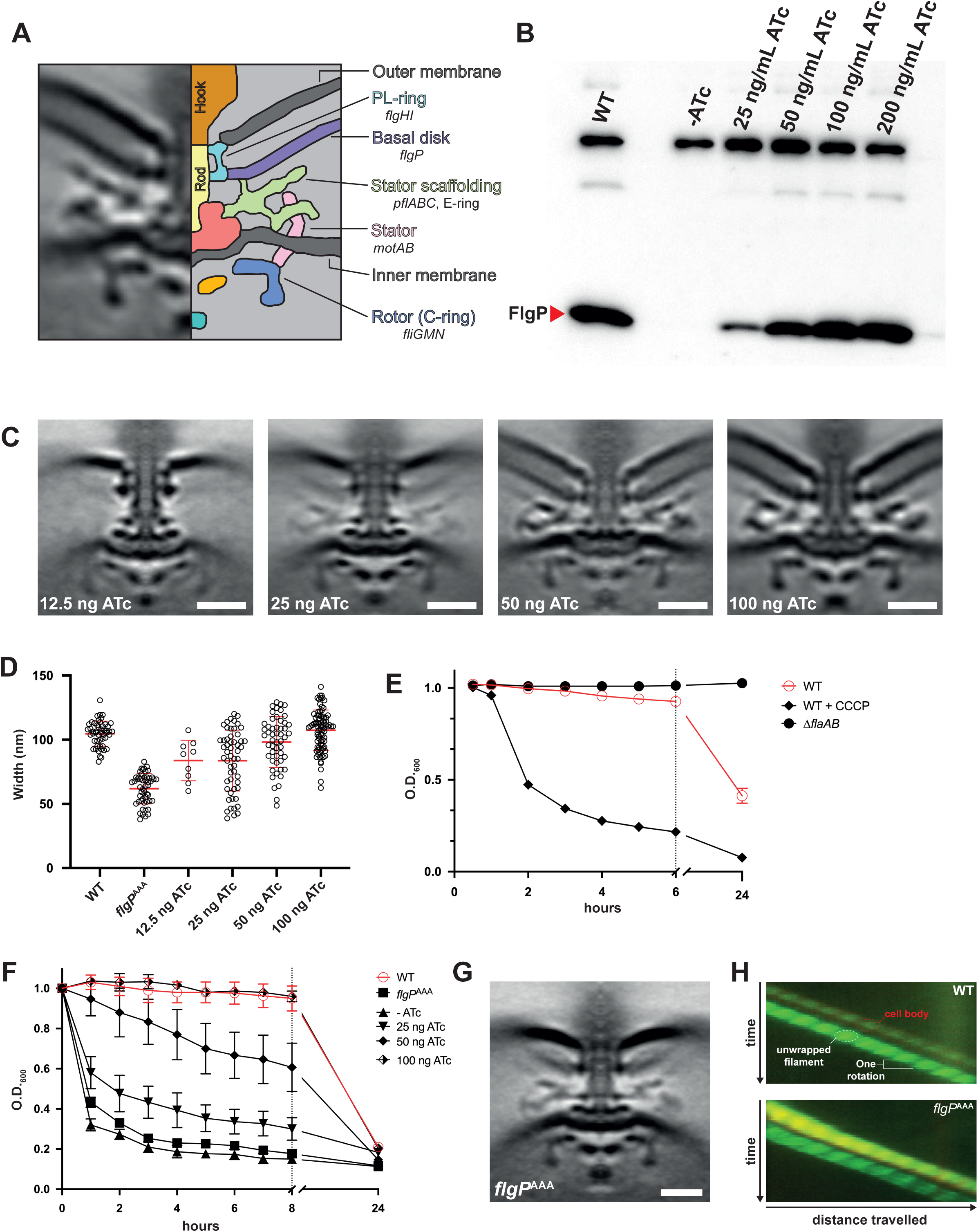
Engineered motors with small basal disks nevertheless incorporate stator complexes and rotate similarly to WT motors. **(A)** The flagellar motor of *Campylobacter jejuni* has the same components as that of the model organisms, as well as extra embellishments such as the basal disk and stator scaffolding. **(B)** Increasing the level of ATc in the growth medium corresponded to an increased level of *flgP* expression and wider basal disks as visualised by **(C)** STA (scale bars = 20 nm) and **(D)** measurements of disks in individual tomograms. In WT, all the motors imaged possessed basal disks, while in EJC168 lower concentrations of ATc corresponded to a lower proportion of motors with disks (12.5 ng ATc/mL: 9% w/ disk; 25 ng ATc/mL: 35% w/ disk ; 50 ng ATc/mL: 75% w/ disk; 100 ng ATc/mL: 88% w/ disk). Disk-less motors were excluded from the analysis in **(D)**. **(E)** Autoagglutination in *C. jejuni* is flagellar-filament dependent, aflagellate mutants do not sediment in autoagglutination assays over the timeframes used in this study. Sedimentation rate is enhanced when the flagellar motor is paralyzed by, *e.g.*, addition of the protonophore carbonyl cyanide *m*-chlorophenyl hydrazone (CCCP) to the cell suspension. **(F)** Sedimentation rate was inversely proportional to *flgPQ* induction level, with non-induced and low induction (25 ng ATc/mL) cell suspensions sedimenting faster than WT and high induction suspensions. **(G)** The *flgP*^S69A^ ^E157A^ ^K159A^ (*flgP*^AAA^) mutant constructed small-diameter basal disks, and the distribution of basal-disk diameters clustered more tightly than EJC168 at all induction levels and never extended beyond ∼80 nm. **(H)** Kymographs of fluorescently-labelled WT and *flgP*^AAA^ cells show that motor-rotation rate is unaffected by small-diameter disks.

We speculated that the large basal disk in *C. jejuni* has functions beyond being the assembly platform for the periplasmic scaffold and wider stator complex ring. We hypothesised that the large diameter of the *C. jejuni* basal disk is an adaptation that allows the disk to act as a flange to stabilise the high-torque *C. jejuni* flagellar motor during unwrapping. Here we present a series of structural, genetic and microscopic experiments consistent with this model.

## Results

### Motors with small basal disks function similarly to motors with large disks

We sought to isolate the contribution of disk diameter from the requirement of the disk for a functional motor. Deletion of the genes responsible for assembly of the basal disk, *flgPQ*, prevents stator complex scaffolding and paralyses the flagellum. To assess the significance of disk diameter, we made a mutant (EJC168) that constructs narrower basal disks but still enables the stator complex periplasmic scaffold by deleting *flgPQ* from chromosome and expressing them *in trans* from a titratable, synthetic *C. jejuni tetRA* promoter system at the *astA* locus^9^. In the absence of inducer anhydrotetracyline (ATc), cells were non-motile) and did not express detectable levels of FlgP. Increasing the concentration of ATc from the lowest concentration tested (0.0125 ug/mL) to the highest tested (0.2 ug/mL) resulted in a corresponding increase of *flgP* expression (Fig. 1B).

Electron cryotomography and subtomogram averaging (STA) of motors from cells grown at different ATc concentrations confirmed that disk size correlated with induction level (Fig. 1C, D). Disks were absent in the absence of ATc, as were stator complexes, equivalent to *flgP* or *flgQ* deletion^8^. Motors assembled at low *flgPQ* expression had narrower disks but were able to assemble the periplasmic scaffold and MotAB stator complexes. Increasing *flgPQ* expression produced correspondingly wider basal disks, with average diameters 84 nm, 98 nm, and 107 nm for 0.025, 0.05, and 0.1 ug/mL ATc respectively. We could discern periplasmic scaffold densities in disks as narrow as 50 nm, whereas the average WT motor is 105 nm, but can polymerize as wide as 130 nm. The WT disk diameter is therefore excessive for a role in stator complex assembly alone.

Consistent with this, motor rotation was not compromised by small disks (Supp. Fig. 1). Across all induction levels, both swimming velocity and filament rotation rate were similar to WT cells, demonstrating that motor function is independent of disk width, provided a minimal disk to facilitate stator complex assembly is present. This implicates the disk in functions beyond rotation.

Despite small-disk motors rotating their flagella at WT speeds, we noticed that suspensions of cells grown on lower concentrations of ATc formed large clumps of cells bound together by their flagellar filaments within minutes of being applied to the sample chamber for observation, unlike suspensions of WT and high-induction cells (Supplemental videos 1- 3). This clumping, known as autoagglutination in *C. jejuni*, is an adaptive behaviour important for microcolony formation during host colonization^10,11^. Autoagglutination requires a flagellar filament, and mutants with impaired or non-functional motors autoagglutinate faster than WT (Fig. 1E). Because small-disk suspensions autoagglutinate faster than large-disk suspensions, despite comparable motor rotation speeds, we conclude that a larger basal disk counteracts autoagglutination (Fig. 1F).

Although disk diameter correlated with level of *flgPQ* induction, we found broad disk diameter distributions (Fig. 1E). We therefore attempted to engineer a mutant that assembles consistently narrow basal disks while remaining motile in soft agar by performing alanine-scanning mutagenesis in *flgP*. We found that *flgP*^S69A^ ^E157A^ ^K159A^ (*flgP*^AAA^) assembled a disk comparable in diameter to that of EJC168 at low induction (0.025 ng/mL ATc), albeit with less size variation (Fig. 1G). Based on western-blot analysis of FlgP, we conclude that FlgP levels in *flgP*^AAA^ are reduced to a fraction of WT through an as-yet unknown mechanism (Supp. Fig. 2). We found that basal disk diameters in *flgP*^AAA^ cluster more tightly than in the ATc induction series and never extend beyond ∼80 nm in diameter (Fig. 1D), but nevertheless rotate their flagella and swim linearly at velocities comparable to WT, demonstrating that the width of the WT basal disk is unnecessary for motor function in isolation (Fig 1H). We found that *flgP*^AAA^ cell suspensions sediment faster than EJC168 induced at 25 ng/mL ATc despite having a higher proportion of cells with disks (44% w/ disks), further supporting that a smaller basal disk fails to counteract excessive autoagglutination.

### The basal disk must be in register with the P-ring for effective motility

That small disks fail to prevent excessive autoagglutination suggested to us that the disk might act as a mechanically-reinforcing flange to stabilise the high-torque *C. jejuni* motor to overcome immediate autoagglutination whenever near another cell. The basal disk is in register with the P-ring, thought to act as a non-rotating bushing to brace the rotating flagellar driveshaft. We reasoned that we could test this model by shifting the basal disk out of register with the P-ring, disrupting mechanical support of the P-ring by the basal disk, yet preserving the disk’s role in stator scaffolding.

We engineered a series of FlgP mutants that pushed the disk out of register with the P-ring. FlgP is a small protein with an N-terminal cysteine (C17), presumed to be lipoylated and inserted in the inner leaflet of the outer membrane (OM). A poorly-conserved N-terminal ∼40 residues of the mature protein is likely to be a linker anchored to the outer membrane by C17. This is supported by a ∼6 nm unresolved gap between the OM and basal disk in the STA structure of the *C. jejuni* motor, a distance consistent with a ∼40-residue linker. To displace the disk, we inserted heptad-repeats of varying length from the *Salmonella* lipoprotein LppA (a.k.a. Braun’s lipoprotein) downstream of C17 (Fig. 2A), followed by the native FlgP OM-linker sequence, based on previous success using Lpp to alter spacing in the periplasm^12–14^. These mutants were motile in soft agar, but motility decreased as more heptads were added (Supp. Fig. 3A and B). We chose the mutant that formed the smallest swarm in soft agar, *flgP-lpp*^55^, for further analysis.

**Figure 2:**
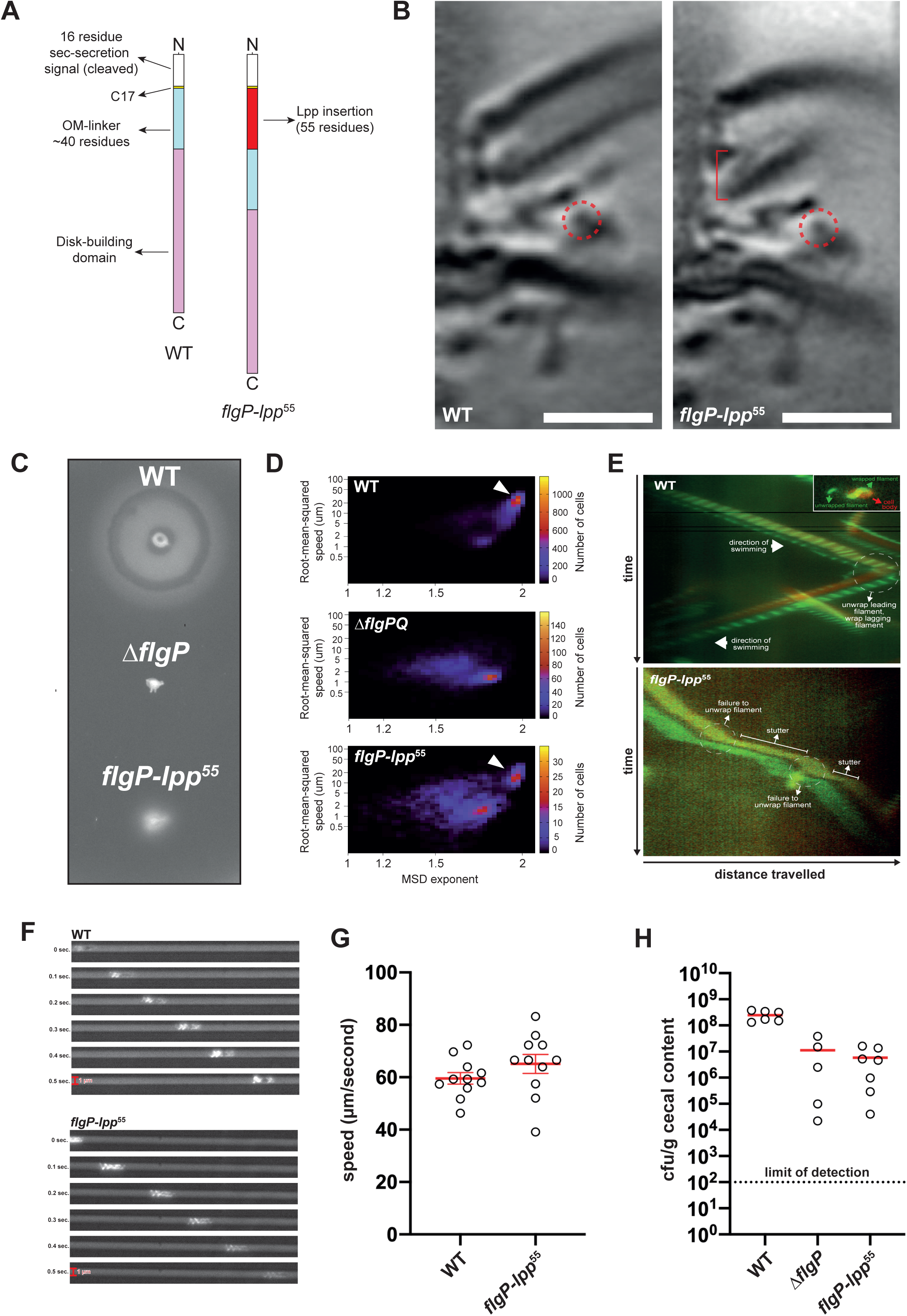
Displacement of the basal disk from the P-ring reduces filament unwrapping. **(A)** A 55-residue segment from *Salmonella* LppA was inserted after C17 in FlgP to make FlgP-Lpp^55^. **(B)** STA of the *flgP-lpp*^55^ motor revealed that the basal disk had been displaced from the P-ring by ∼7 nm (red bracket), but that stator recruitment was not impacted (dashed red circles)(scale bars = 20 nm). **(C)** The *flgP-lpp*^55^ mutant formed small-diameter swarms in motility agar compared to WT, despite **(D)** the presence of a population of cells swimming at WT velocity when cell suspensions were observed by 3D-holographic-tracking microscopy (arrowheads). **(E)** High-speed-fluorescence-video microscopy showed that the *flgP-lpp*^55^ mutant was incapable of unwrapping its filament from the cell body during motor reversals, resulting in a stuttering motility rather than the characteristic darting motility of WT *C. jejuni*. **(F** and **G)** The velocity of individual *flgP-lpp*^55^ cells traversing a microfluidic device with 1 μm- wide channels was found to be identical to WT, confirming our observations from holographic tracking. **(H)** The *flgP-lpp*^55^ mutant colonizes the chicken cecum as poorly as a non-motile Δ*flgP* mutant.

STA confirmed that the *flgP-lpp*^55^ motor had a basal disk that had been shifted out of register with the P-ring by ∼7-8 nm, consistent with the 8.4 nm expected by insertion of a 56-residue α-helix (Fig. 2B). In this mutant, the disk encircles the proximal rod instead of the P-ring. Curiously, the inner radius of the *flgP-lpp*^55^ disk was narrower than the WT disk, matching the decreased width of the structure around which it was assembling. This supports that the disk self-assembles non-specifically around whichever axial component it is in register with. Although *flgP-lpp*^55^ forms small-diameter swarms in soft agar (Fig. 2C), the stator scaffolding architecture in the *flgP-lpp*^55^ motor was indistinguishable from the WT motor, and 3-D holographic tracking microscopy of the *flgP-lpp*^55^ mutant swimming in viscous media revealed a population of the cells swimming at WT velocity (Fig. 2D).

To determine how displacement of the basal disk from the P-ring reduced motility in soft agar despite the presence of motile cells in liquid media, we labelled the flagellar filament with fluorescent dye and recorded *flgP-lpp*^55^ swimming by high-speed-video fluorescence microscopy. As with our small disk mutants, and consistent with our holographic tracking, flagellar rotation rate in the *flgP-lpp*^55^ mutant was the same as WT, but the mutant was incapable of unwrapping the leading flagellum from the cell body during motor reversals (Supplemental video 4). Consequently, *flgP-lpp*^55^ exhibited a stuttering motility characterised by short runs interrupted by brief pauses, with no net change in swimming trajectory (Fig. 2E). This contrasts with WT cells, which have a swimming style referred to as darting motility, whereby chemotaxing cells change swimming trajectory during motor reversals due to the wrapping and unwrapping of the opposed flagellar filaments (Supplemental video 5). Thus, the phenotype of the *flgP-lpp*^55^ mutant in soft agar reflects the inability of the filament to be pulled from the cell body during motor switching across the entire population. Using darkfield microscopy, we found that the failure of *flgP-lpp*^55^ to unwrap upon motor reversal results in population-level failure to swarm toward regions of higher oxygen content in aerotaxis assays, unlike WT populations (Supplemental video 6).

To understand how this impairment would affect motility in environments like those of the mucous-filled gastric crypts of a host’s digestive tract, we observed fluorescently-labelled cells swimming through viscous media in a microfluidic device with confined 1 μm channels, allowing only one cell through at a time (Supplemental video 7). While individual *flgP-lpp*^55^ cells were able to traverse the device at the same speed as WT cells (Fig. 2F and G), further confirming our STA results and 3D holographic cell tracking, the displaced-disk *flgP-lpp*^55^ mutant was unable to reverse direction when encountering obstacles (*i.e.* immobilized cells) in the channel, leading to *C. jejuni* logjams (Supplemental videos 8 and 9). In contrast, WT cells reversed direction upon encountering an obstacle and exited the device (Supplemental video 10). We next inoculated day-old chicks with identical numbers of WT, Δ*flgP* or *flgP- lpp*^55^ cells and enumerated the number of colony forming units (cfu) in the ceca of the chicks at one week post-inoculation. As predicted by our microfluidic device results, we found that the *flgP-lpp*^55^ mutant colonized the chicken cecum as poorly as the non-motile Δ*flgP* control strain. These results show that, in addition to the basal disk’s role in stator scaffolding, flanging of the motor by the basal disk is an important adaptive trait for *C. jejuni* in its native environments.

We also produced a mutant in which the disk was moved closer to the outer membrane, and out of register with the P-ring, by deletion of the OM-linker while retaining C17 (*flgP*^Δ18–62^) (Supp. Fig. 3C and D). In soft agar, *flgP*^Δ18–62^ produced a swarm ∼30% that of WT. In contrast to *flgP-lpp*^55^, however, the subtomogram average of the *flgP*^Δ18–62^ motor had poorly resolved stator scaffolds, suggesting that the motility defect in this background is at least partially due to low average stator occupancy in the motor, and consistent with FlgP being too far from the inner membrane to template formation of the periplasmic scaffold (Supp. Fig. 3E).

In sum, these results show that a wide disk is required for overcoming filament-filament interactions during autoagglutination, while a correctly-positioned disk is required to act as a flange to overcome filament-cell interactions during unwrapping.

### Mutations that restore motility to a displaced disk mutant suggest a link between filament unwrapping and filament glycosylation

Suppressing mutations in the *flgP-lpp*^55^ background that restored near-WT levels of motility arise following 36-48 hours of incubation, which appear as flares emanating from the original, poor-motility swarm. Reasoning that the identity of the suppressor mutations would be further informative as to the function of the basal disk, we isolated two independent revertants, performed whole genome sequencing, and identified two paths to suppress the *flgP-lpp*^55^ motility defect: restoration of the P-ring/basal disk register (*i.e.* restoration of flanging), or decreased *O*-glycosylation of the flagellar filament.

The first motile revertant of *flgP-lpp*^55^ acquired a *flgG\** mutation, *flgG*^T54N^. In WT flagellar motors, the distal rod (FlgG) only extends enough for a single P-ring and a single L-ring before it contacts the outer membrane and stops polymerizing. Alleles in *flgG*, however, have been isolated that allow it to continue polymerizing^15^. These so-called *flgG\** alleles often arise in an N-terminal region of FlgG known as the Dc domain, encompassing residues ∼30-70 of the protein^16–18^. Similarly, our *flgG*^T54N^ allele allows the distal rod to grow longer than usual to accommodate two P-rings around the distal rod (Fig. 3A); thus the additional space imposed by the Lpp^55^ insertion is compensated for by an additional P-ring on a flgG^T54N^ rod, thus restoring register of the first P-ring to the basal disk, while the second P- ring templates assembly of the L-ring for correct OM-penetration and hook/filament assembly.

**Figure 3:**
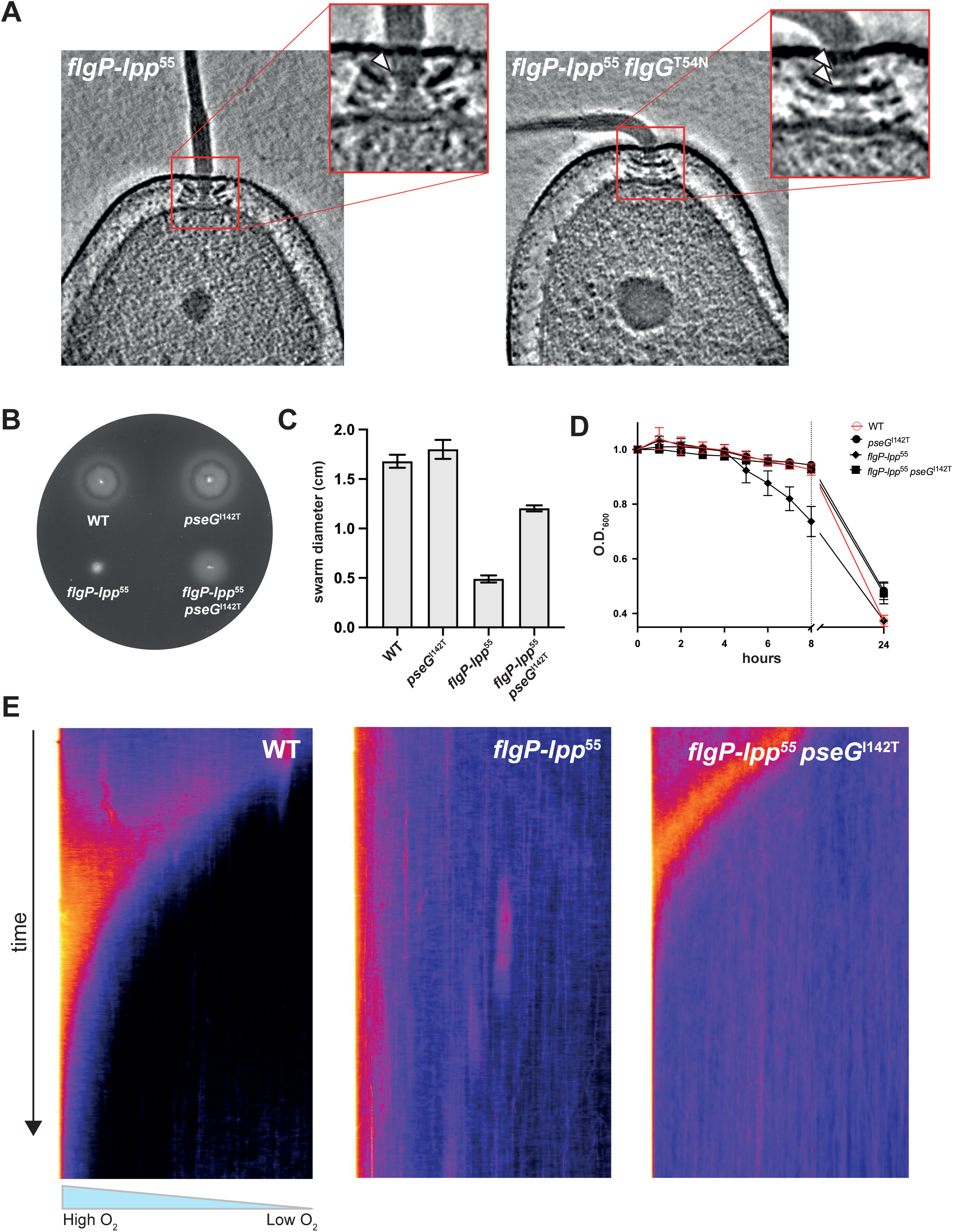
Suppression of the *flgP-lpp*^55^ motility defect occurs by restoring the basal disk-P-ring register or reducing filament glycosylation. (A) The *flgG*^T54N^ allele allows the distal rod to grow longer in order to accommodate a second P-ring (arrowheads), restoring flanging of the motor by the basal disk in the *flgP-lpp*^55^ background. **(B and C)** The second suppressor we isolated arose in *pseG*. The *pseG*^I142T^ allele has no effect on motility in an otherwise WT background but increases soft-agar motility of the *flgP-lpp*^55^ mutant. **(D)** The decreased sedimentation rate caused by the *pseG*^I142T^ allele in both the WT and *flgP-lpp*^55^ backgrounds indicates lower levels of *O*- glycosylation of the filament. **(E)** The *pseG*^I142T^ allele allows unwrapping of the filament from the cell body upon motor reversal in the *flgP-lpp*^55^ background, demonstrated at the population level by restored aerotactic behaviour of the *flgP-lpp*^55^ *pseG*^I142T^ double mutant.

The second revertant was intriguing because the suppressing mutation, *pseG*^I142T^, affects the flagellar filament rather than the motor. PseG is a UDP-sugar hydrolase involved in the synthesis of pseudaminic acid (PseAc)^19,20^, the *O*-linked sugar that decorates the flagellar filament at 19 residues of each flagellin monomer in *C. jejuni* 81-176^21–23^. Glycosylation of the flagellar filament in *C. jejuni* is required for both filament assembly as well as autoagglutination. Only three of the 19 glycosylable flagellin residues are critical for filament assembly and motility, and a *pseG* knockout is non-motile^24,25^. We therefore reasoned that substitution of a non-polar isoleucine residue adjacent to the substrate-binding site of PseG with a threonine, decreases, but must not abolish, its enzymatic activity and results in filaments with reduced *O*-glycosylation. This interpretation is supported by the *pseG*^I142^ allele in an otherwise WT background having no effect on swarm diameter in soft agar (Fig. 3B and C). Furthermore, the *pseG*^I142T^ allele reduces autoagglutination in both the WT and *flgP- lpp*^56^ backgrounds (Fig. 3D), in agreement with previous work showing that eight of 19 glycosylable flagellin residues are important for autoagglutination^21^.

When we observed the *flgP-lpp*^55^ *pseG*^I142T^ double mutant by high-speed-video fluorescence microscopy, we found that the double mutant unwrapped its filament from the cell body during directional reversals at rates comparable to WT, in contrast to *flgP-lpp*^55^. Furthermore, in both aerotaxis assays and microfluidic experiments the *flgP-lpp*^55^ *pseG*^I142^ mutant exhibited near-WT behaviour (Fig. 3E). Consistent with our previous study showing interaction between the flagellar filament and cell body, these data implicate *O*-glycosylation of the flagellar filament in filament-cell body interactions in addition to filament-filament autoagglutination and filament assembly.

### Swimming without a basal disk requires de-glycosylation of the cell surface

Our results indicate that a small disk is sufficient for motor function, albeit insufficient to counteract autoagglutination. Although deletion of *flgPQ* produces a non-motile phenotype in motility agar, observing this mutant by fluorescence microscopy revealed occasional cells with rotating flagella. Thus, even in the absence of the basal disk, stator complexes can still be inefficiently recruited to the motor, although those Δ*flgPQ* cells with rotating flagella tend to be found in clumps of autoagglutinated cells. Given that we were able to isolate suppressors of disk displacement, we speculated that we might be able to isolate a suppressor strain of a wholesale *flgPQ* deletion that would shed more light on the role of the basal disk.

We selected for suppression of the Δ*flgPQ* motility defect through prolonged incubation in motility agar, as previous attempts over smaller time frames (∼48-72 hours) had been unsuccessful in isolating motile revertants. We independently inoculated two colonies each of Δ*flgPQ* and Δ*flgQ* mutants in motility agar and incubated the plates for four to six days. Although each colony had a non-motile phenotype after two days of incubation, all four isolates had speckles emanating from the site of inoculation within five days.

The speckled phenotype occurs when the majority of cells in motility agar are non-motile, with occasional cells possessing a functional flagellum. These motile cells deposit non-motile daughter cells upon division which seed colonies of non-motile descendants. As this process occurs around the site of inoculation, the swarm takes on a speckled, or “bushy”, phenotype. After four to six days of incubation, the edge of each bushy swarm was picked from the agar, single-colony purified, stored at -80C, and also used to inoculate a fresh motility plate. This process was repeated four to five times for each lineage, at which point all lineages had evolved a smooth-swimming motility-swarm phenotype (Fig. 4A), indicating that a majority of cells in the population are swimming. We then performed whole-genome sequencing of each endpoint isolate for each of the four lineages to determine the mutations required for motility in the absence of the basal disk.

**Figure 4:**
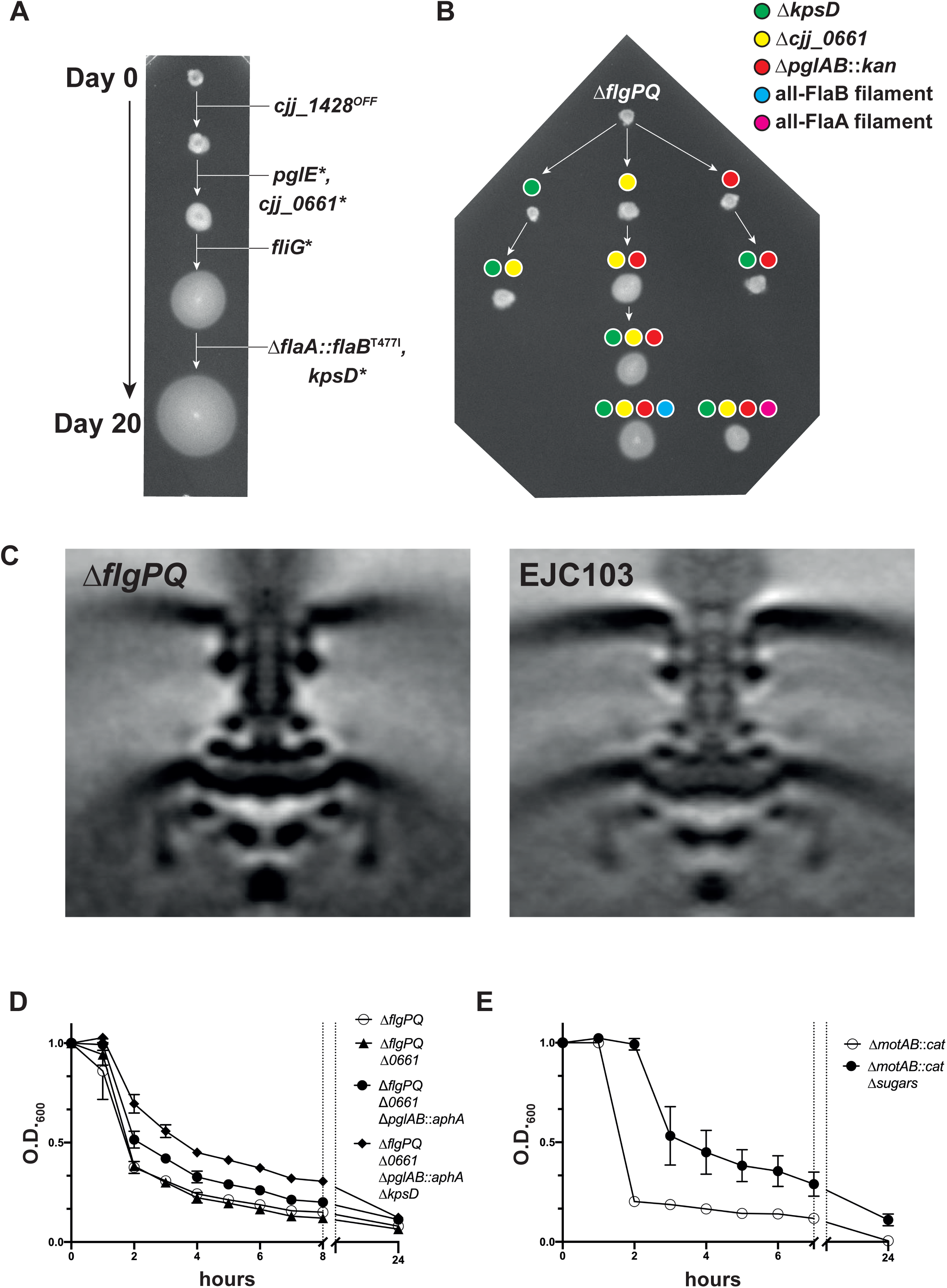
Motility in the absence of a basal disk requires deglycosylation of the cell surface. **(A)** Four independent disk-less cultures were inoculated into soft agar and allowed to incubate several days. Speckles emanating from the point of inoculation were picked, purified, stored and also used to inoculate a subsequent incubation in soft agar. After repeating four to five times for each independent lineage, all lineages had evolved a “smooth swimming” phenotype. **(B)** Generating *pgl*, *kps* and *0661* knockouts singly and in combination revealed that the spontaneous mutations from the evolution experiment were loss-of-function mutations and that loss of both *0661* and *N*-glycosylation (*i.e. pgl*) are required to suppress the Δ*flgPQ* motility defect **(C)** Comparison of the STA structures of the Δ*flgPQ* and EJC103 mutants’ motors confirmed that motility in the evolved disk-less mutants was not due restoration of stator scaffolding in the motor. **(D)** Suppression of the non-motile phenotype in the absence of the basal disk corresponded with decreased sedimentation rate upon removal of *pgl* and *kps* surface polysaccharides in autoagglutination assays. **(E)** Comparison of the sedimentation rates of cells paralyzed by deletion of the stators (Δ*motAB*::*cat*) in both the WT background and the Δ*0661* Δ*pglAB*::*aphA* Δ*kpsD* background shows that the decreased sedimentation rate of the of EJC103 in **(D)** is not a result of increased motility in this background.

We predicted that disk-less motility would require mutations in flagellar genes, specifically periplasmic scaffold genes and/or the stator complex genes *motAB*, as such mutations might enable stable incorporation of stator complexes despite lack of the scaffolding role of FlgP. To our surprise, however, the theme across all four evolved lineages was a similar constellation of mutations in genes involved, or implicated, in decorating the cell surface with polysaccharides (Table 1). Each lineage had mutations in the *pgl* operon, responsible for N-glycosylation of a diverse cohort of periplasmic and surface-exposed proteins^26–29^. A further two lineages had acquired mutations in *kps* genes, which are responsible for capsular polysaccharide (CPS) biogenesis^30^. Additionally, mutations in a gene predicted to function as a polysaccharide deacetylase, *cjj_81-176_0661* (hereafter referred to as *0661*), were present in all four evolved lineages. Each lineage also had evidence of phase variation in *kps*-associated sugar transferase and CPS-modification genes^31,32^.

**Table 1:**
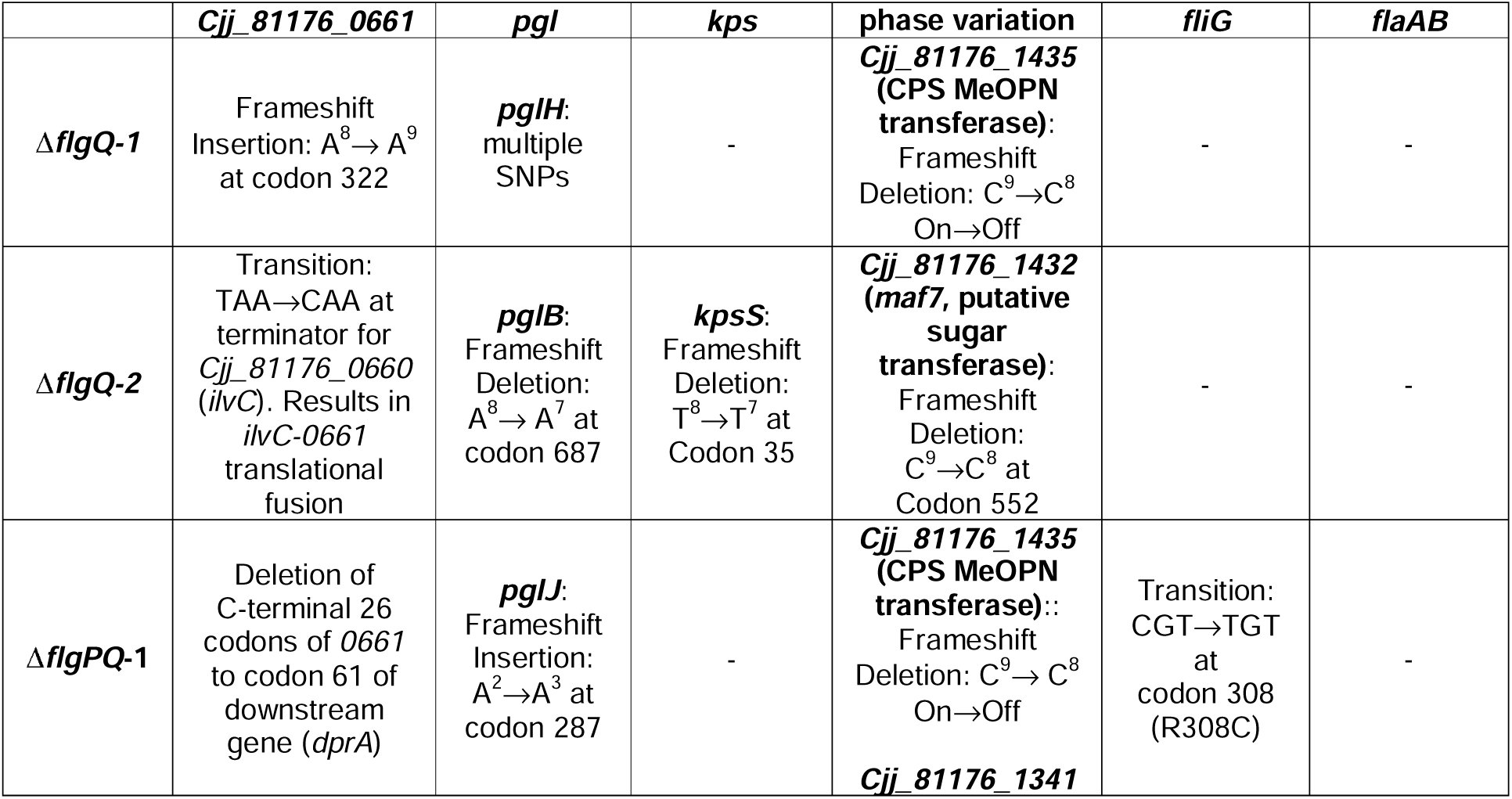
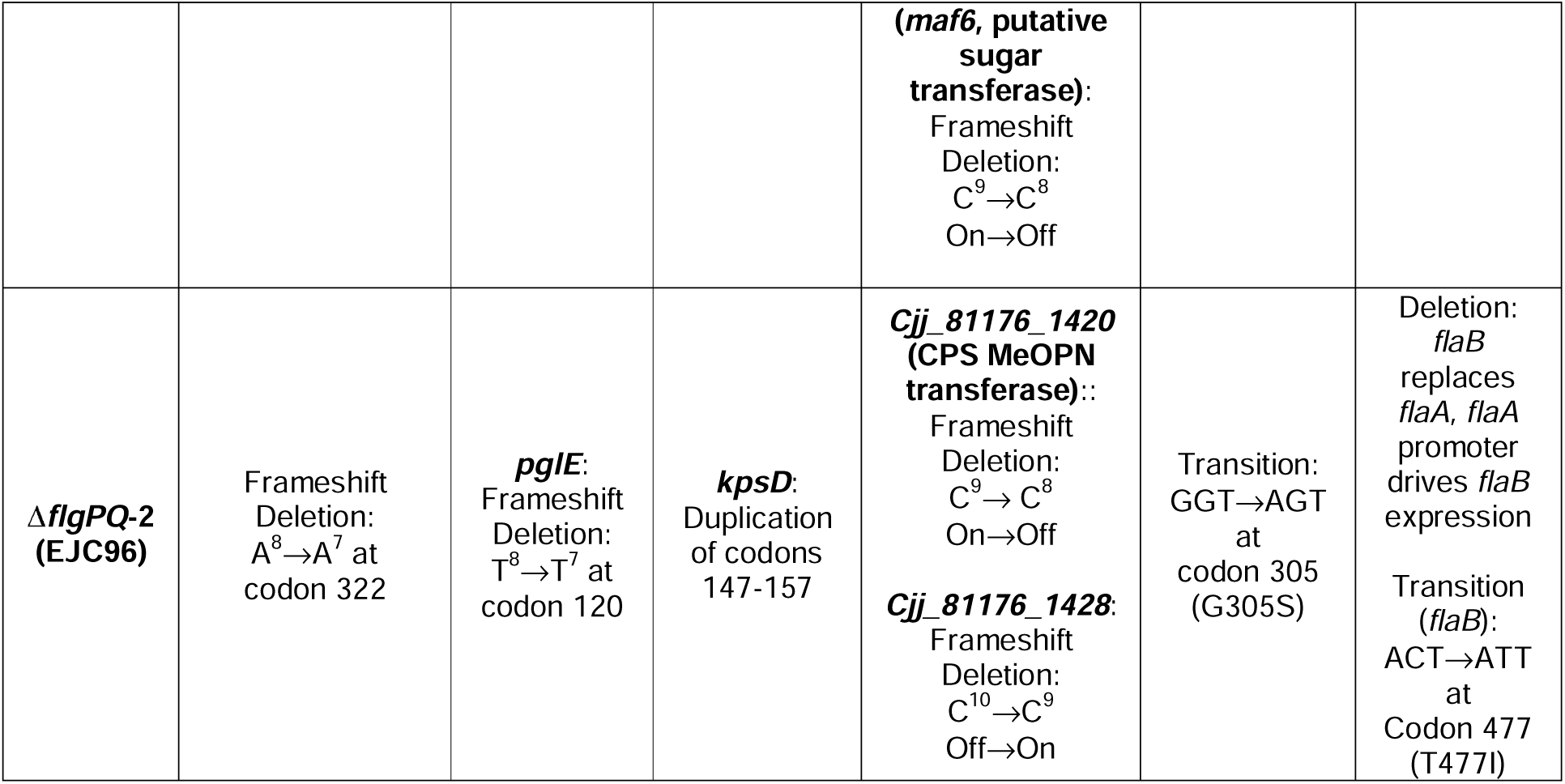
Selection for suppression of theΔflgP(Q) motility defect invariably returned inactivating mutations in cell-surface glycosylation genes.

**Table 2:**
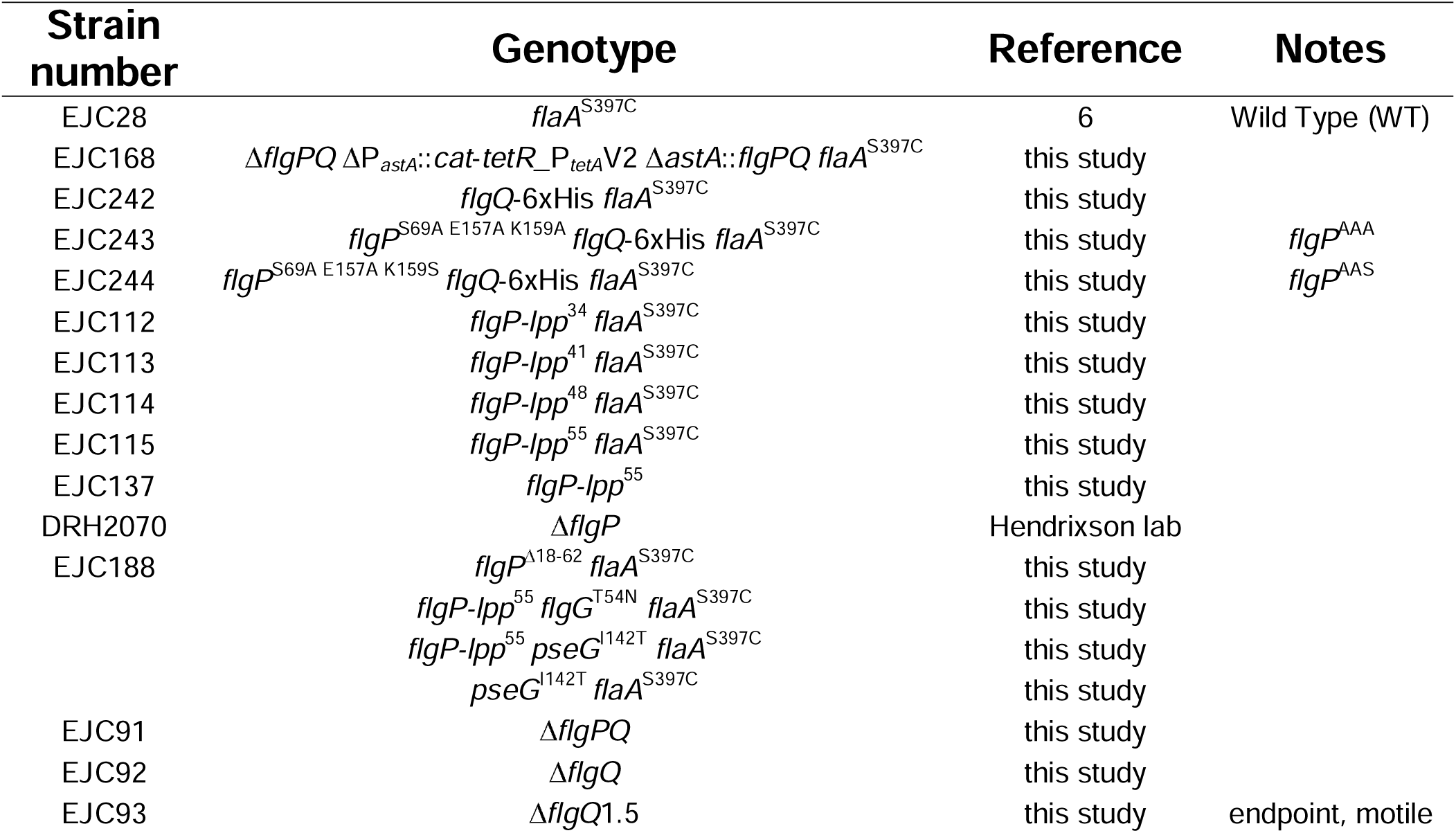
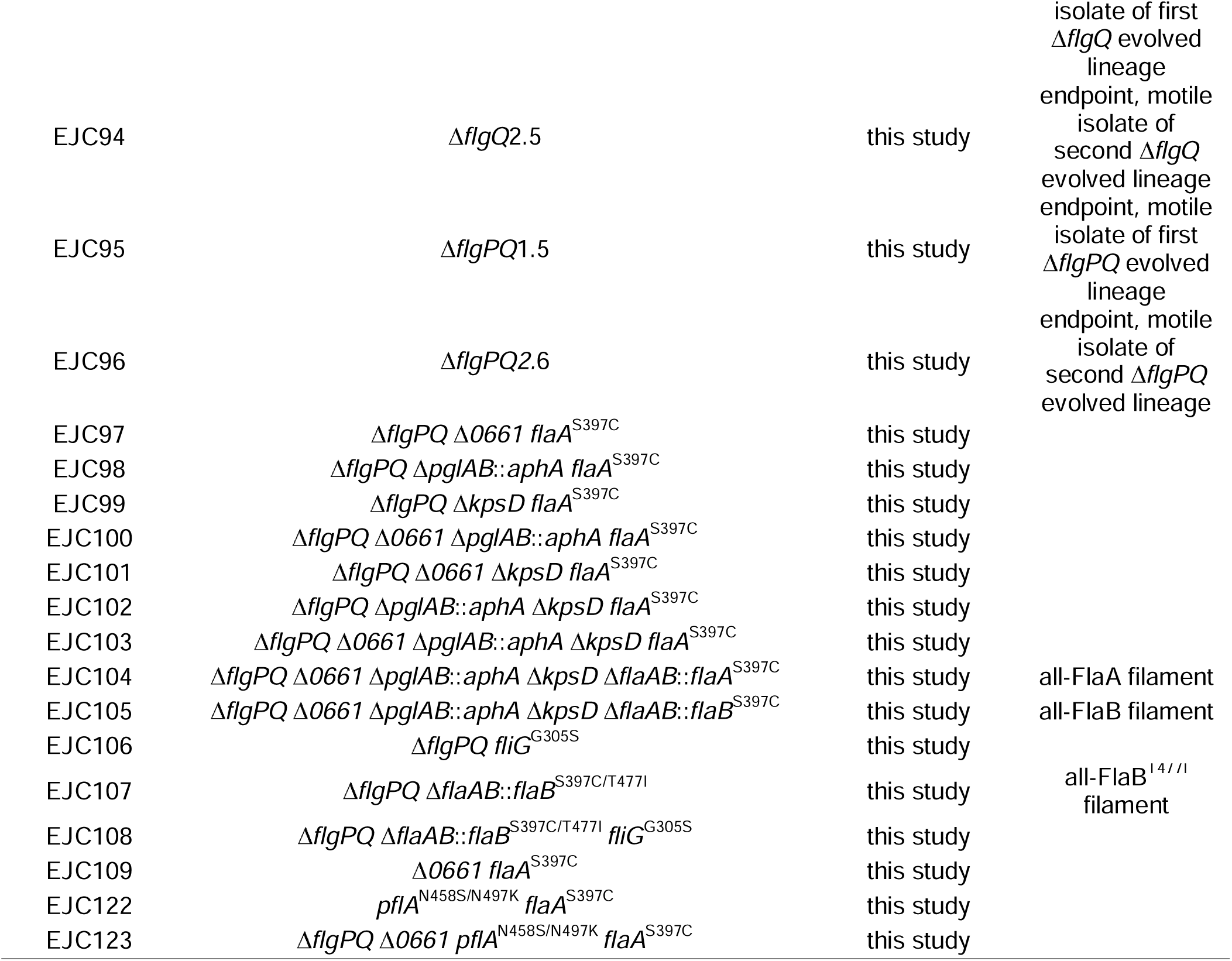
List of strains used in this study.

In addition to mutations in genes involved or implicated in decoration of the cell surface with sugars, two of the isolates had single-nucleotide polymorphisms (SNPs) in the gene encoding the cytoplasmic rotor, *fliG*^33^, and one isolate had a second flagellar mutation at the *fla* locus: Δ*flaA::flaB*^T477I^, in which the the WT flagellin locus encoding both the major flagellin *flaA* and the minor flagellin *flaB* collapsed into one *flaB*-type gene expressed from the *flaA* promoter. To determine the impact of each mutation identified by whole genome sequencing, we selected the endpoint, motile isolate from one lineage, EJC96, for further study as it possessed mutations in *pgl*, *0661* and *kps* genes of as well as the *fliG*^G305S^ and Δ*flaA::flaB*^T457I^ alleles (Table 1).

We sequenced each stored time point of the Δ*flgPQ-2* lineage to determine the order in which each suppressing allele arose. We found that loss-of-function mutations arose in *0661* and *pglE* first, followed by the *fliG*G^305S^, Δ*flaA::flaB*^T457I^ and *kpsD\** alleles (Fig. 4A). To rule out the possibility that the mutations in EJC96 that affect glycosylation were gain-of- function mutations, *i.e.* to confirm that the suppressing effect of the mutations on motility in soft agar was due to the loss of glycosylation, we constructed strain EJC103 (Δ*pglAB*::*aphA* Δ*kpsD* Δ*0661*) which lacks CPS and the ability to *N*-glycosylate proteins as well as *0661*. We found that EJC103 had a smooth-swimming phenotype in soft agar, confirming that suppression of the Δ*flgPQ* motility defect occurs via loss of glycosylation. By constructing deletions of *pglAB*, *kpsD* and *0661* in all combinations (EJC97-103, Fig. 4B), we discovered that loss of *0661* is critical for soft-agar motility in the Δ*flgPQ* background, as the *pglAB*::*aphA* and Δ*kpsD* alleles, either alone or in tandem, are insufficient to promote swimming in the absence of the basal disk.

To assess the importance of the *fliG*^G305S^ and Δ*flaA::flaB*^T477I^ mutations relative to the glycosylation genes, we reconstituted the *fliG*^G305S^ and Δ*flaA::flaB*^T477I^ mutations singly and in combination in the Δ*flgPQ* parental background. We found that neither *fliG*^G305S^ nor Δ*flaA::flaB*^T477I^ alone suppress the Δ*flgPQ* motility defect, while the double mutant displays some degree of speckling but is otherwise indistinguishable to the Δ*flgPQ* parental strain (not shown). These results show that loss of cell-surface glycosylation is necessary and sufficient to permit swimming in the absence of a basal disk, while flagellar-specific mutations only serve to enhance motility once the cell surface has been de-glycosylated.

To investigate the possibility that loss of *pglAB*, *kpsD* and *0661* had somehow restored stator scaffolding in the absence of the basal disk, we performed subtomogram averaging of the Δ*flgPQ* and EJC103 motors and found them to be indistinguishable from one another, with no visible stator complex densities in either, (Fig. 4C) as seen with subtomogram average structures of the *Salmonella* and *E. coli* motors that lack statically-assembled stator complexes.

Indeed, the auto-agglutination profiles from the Δ*flgPQ* parental through to EJC103 revealed that the evolution from immotile to smooth swimming in motility agar coincided with a decrease in autoagglutination rate (Fig. 4D). Taken together, this indicates that in the absence of *flgPQ, C. jejuni* stator complexes can still transiently incorporate into the motor to generate torque, but that motility in WT cells is halted due to immediate autoagglutination. Short of the highly improbable *de novo* evolution of a replacement for *flgPQ,* the only way to restore motilty is via reduction of autoagglutination by disabling cell-surface glycosylation, thereby allowing motile Δ*flgPQ* cells to avoid or escape cell aggregates and remain free-swimming

## Discussion

We sought to understand the selective advantage of the large basal disk in *C. jejuni*. Our findings reveal that the complex architecture of the *C. jejuni* flagellar motor has roles in addition to simply increasing the torque of the motor and ensuring high-stator occupancy.

Taken together, our results implicate the basal disk as a flange that stabilises the motor, enabling the motor to function despite the filament-filament and filament-cell attractions involved in autoagglutination and flagellar wrapping around the cell body, respectively. Our small-disk mutants indicate that a small disk is sufficient to assemble a functional motor whose rotary function is unaffected. The large diameter and proper placement of the wild-type disk, while not necessary for motor rotation, however, is critical for wrenching filaments from one another and from the cell body. We propose that this wrenching requires that the PL-ring bushing around the rod be stabilized more than it is in the model organisms *E. coli* and *Salmonella*. This is graphically illustrated by poor unwrapping when the disk is displaced from the P-ring (by insertion of Lpp in FlgP), and restored unwrapping when a P-ring is placed back in register with the disk (by a *flgG*\* suppressor mutation). The motility impairment arising from small disks or displaced disks can be countered by attenuating sugar-mediated interactions, with both mutants in *O-*glycosylation pathways of the flagellar filament and *N-*glycosylation pathways of the cell surface restoring motility in disk mutants.

*C. jejuni* coordinates its opposing flagella at each pole by wrapping the leading filament around the cell body. In WT cells, directional reversals for chemotaxis occur when motor rotation switches transiently to clockwise, leading to switching which filament is wrapped around the cell body. This requires that the wrapped filament is pulled free from the cell body. What was unexpected in our previous study was the finding that a straight-cell-body mutant struggles to unwrap its wrapped filament. There, we hypothesized that the glycosylated filament and glycosylated cell surface have an affinity for one another, and that helical cell shape minimizes contact between the filament and cell surface.

In this study, we find further support for this hypothesis. Selection for suppression of motility defects in Δ*flgPQ* mutants invariably returned mutations in cell-surface-glycosylation genes, while rare flagellar-specific mutations are neither necessary nor sufficient to promote swimming in the absence of a basal disk. Furthermore, the swimming defects of small-disk and displaced-disk mutants are not due to impairment of motor function, as both possess motors that rotate comparably to WT motors.

Our isolation of the Δ*flaA::flaB*^T477I^ mutation in the EJC96 lineage, in which FlaA is replaced by FlaB, may be explained by our previous finding that an entirely-FlaB filament is more rigid than an entirely-FlaA filament. The EJC96 lineage may have evolved a more rigid filament that is more easily pulled from the cell surface during unwrapping attempts. Indeed, an all- *flaA* filament in the EJC96 background is worse at swimming in motility agar than either *flaA*^+^*B*^+^ or all-*flaB* in the same background, with all-*flaB* being the best of the three.

Inevitably, our results have limitations and unresolved complexities. One of the principal complications is that our suppression experiments returned a combination of mutations affecting different types of surface-sugar interactions: filament-filament interactions involved in autoagglutination and filament-cell body interactions involved in unwrapping during directional switching. Curiously, the suppressors of different types of mutant differ: disk displacement (in the *flgP-lpp*^56^ mutant) is suppressed by reducing filament *O*-glycosylation, while disk removal (by deleting *flgPQ*) is suppressed by abolishing cell surface glycosylation (by preventing capsule biosynthesis and N-linked glycosylation). Introduction of the *pseG*^I142T^ allele into the EJC103 background background has no noticeable effect on motility in soft agar. Similarly, a *flgP-lpp*^55^ Δ*pglAB*::*aphA* Δ*kpsD* Δ*0661* mutant swims no better than *flgP- lpp*^55^ alone

Why do we only see cell surface glycosylation suppressors, and not filament glycosylation suppressors of disk-less mutants? We speculate that the *pseG*^I142T^ allele fails to further boost motility in the EJC103 background (lacking capsular polysaccharide, the ability to N- glycosylate proteins, and enigmatic protein 0661) mutant because the *pseG*^I142T^ strain’s filament remains heavily glycosylated, and a low torque (due to low-stator-occupancy) Δ*flgPQ* motor cannot overcome the still-strong affinity between two glycosylated filaments. That the difference in autoagglutination profiles between WT and *pseG*^I142T^ are only apparent following extended incubation periods, in contrast to WT vs. Δ*flgPQ*, suggests that the effect of the *pseG*^I142T^ allele on filament glycosylation is small, and therefore beneficial to a mutant whose motor is only moderately impaired. Indeed, the *flgP-lpp*^55^ mutant autoagglutinates only somewhat faster than WT, a defect that is rescued by the *pseG*^I142T^ allele. This suggests that although flanging in the *flgP-lpp*^55^ motor is disrupted, the otherwise-functional motor in this background remains sufficiently powerful to overcome filament-filament adhesion.

Similarly, we found that deletion of surface glycosylation genes (*i.e. pglAB* and *kpsD*) in the *flgP-lpp*^55^ background does not suppress its motility defect in soft agar (not shown). Why is this the case? Our results suggest that the defects in soft-agar motility in the Δ*flgPQ* background and *flgP-lpp*^55^ background are fundamentally different. The motility defect in the Δ*flgPQ* background is due not only to inefficient stator incorporation into the motor, but also enhanced autoagglutination. Loss of cell surface sugars, therefore, allows Δ*flgPQ* cells to escape cell aggregates and remain free swimming. In contrast, the *flgP-lpp*^55^ defect appears to be almost entirely due to its unwrapping defect, as it has only a somewhat enhanced autoagglutination rate. Thus, mutations that allow diskless cells to avoid cell clumps are not expected to alleviate the motility defect of the *flgP-lpp*^55^ mutant. A remaining mystery is what specifically the *O*-glycosylated filament is interacting with on the cell surface, as it does not appear to be either surface-exposed *N*-glycosylated proteins or CPS. We speculate that the filament may have an affinity for lipooligosaccharide or another surface-exposed moiety, but this will require further investigation.

Another factor that may contribute to the relatively small autoagglutination defect of *flgP- lpp*^56^ is that filaments which remain wrapped around the cell body are not available to participate in the filament-filament adhesion that is the basis of autoagglutination, *i.e.* only one half of the filaments in a suspension of cells are unwrapped and thus free for autoagglutination. This may help to explain why small-disk motors have pronounced autoagglutination defects, even though the *flgP-lpp*^55^ motor constructs basal disks with an average diameter of ∼70 nm, smaller than disks of EJC168 grown on 50 ng/mL ATc. We were surprised that we didn’t observe an obvious unwrapping defect in small-disk motors; it is possible that unwrapping doesn’t require an extra-large basal disk so long as the disk is in register with the P-ring.

The role of *0661*, deletion of which is required to suppress the Δ*flgPQ* motility defect, remains unclear. The gene product of *0661* is a member of the PF04748 divergent polysaccharide deacetylase family and is ubiquitous across *Campylobacterota*, but is absent from other genera (Supp. Fig. 4). All Campylobacterota *0661* homologues form a discrete clade within the PF04748 phylogeny, indicating that they retain a common function.

Curiously, PF04748 occurs even in the absence of capsule or flagellin glycosylation genes, suggesting an alternative role. The ubiquity of *0661* in the *Campylobacterota* indicates that this gene is ancient and arose shortly after the *Campylobacterota* genus branched off from other bacterial taxa. While we observed no impact of deleting *0661* on growth and motility in an otherwise WT background in a laboratory setting, *0661* is essential for colonization of the chicken cecum^34^.

We speculate that *0661* may process peptidoglycan based on three observations: 1) *0661* is predicted to be an inner-membrane-anchored periplasmic protein, similar to another peptidoglycan acetyltransferase, PatB, in *C. jejuni*^35^, 2) deletion of *0661* has no effect on autoagglutination rate in the Δ*flgPQ* background, and (3) deletion of *0661* in the Δ*flgPQ* mutant results in a transition from ∼5% of cells rotating their flagella to ∼25% of cells having rotating filaments (Supplemental video 11), while there is no apparent further increase as *pglAB* and *kpsD* are knocked out, suggesting deletion of *0661* somehow enhances the ability of stators to associate with the motor in the absence of scaffolding.

In the model organisms, the stator protein MotB is believed to bind peptidoglycan via a catch-bond mechanism and has been shown to co-crystallize with *N*-acetylmuramic acid, a component of peptidoglycan^36^. It is possible that *0661*’s enzymatic activity defaces a binding motif on peptidoglycan required for efficient recruitment of *Camplyobacterota* MotB to the motor, which has been rendered unnecessary for motility in the *Campylobacterota* due to the ubiquity of stator scaffolding across this genus.

Despite the uncertainties and complexities in our findings, the common theme that runs through all of our findings is the interconnectedness of surface-sugar-mediated interactions and a flanged, high-torque flagellar motor in *C. jejuni*.

Investigation of the myriad functions of the cell-surface glycome of *C. jejuni* has been an area of intense research for several decades^37,38^; *N*-linked glycosylation has been been shown to be important for protein folding, protection of proteins from proteolytic degradation, natural transformation, and adhesion to host cells^39–43;^ CPS is known to protect against insult by antimicrobials and bacteriophage in the environment, to help modulate and evade the host immune system, and is involved in biofilm formation and host colonization^31,44,45^; *O*- glycosylation of the flagellar filament, in addition to promoting filament assembly and autoagglutination, is known to modulate adhesion to host epithelial cells^46,47^. Overall, *C. jejuni* devotes a relatively large proportion of its genome (>8%) to genes involved in the decoration of the cell surface with polysaccharides^38^. As *C. jejuni* has evolved an ever more complex and abundant collection of surface-exposed sugars that it uses to thrive in its environmental niche it has become, in a word, sticky (which appears to be, in many instances, “the point”).

We posit that *C. jejuni* has glycosylated itself into an evolutionary corner: the once- dispensable basal disk and associated stator scaffolding have become indispensable as the cell has become more and more sticky. The structural complexity of the *C. jejuni* motor, however, can be reduced to that of a *Salmonella*-type flagellar motor and promote swimming, provided devolution of cell-surface polysaccharides occurs in tandem. In addition to allowing free-swimming and unwrapping by individual planktonic cells, we speculate that a flanged motor is important for the dispersal of individual cells from a sessile biofilm glued together by filaments and surface sugars as cells seek out new sites for colonization in their environment.

The most important outstanding question to be addressed is the mechanism by which the disk stabilises the motor to enable it to overcome the attractive forces between filaments and cells. Of considerable confusion to us, stabilisation of the motor by the basal disk does not appear to depend on anchoring the basal disk in the OM via lipidation of FlgP’s N-terminal cysteine. A *flgP*^C17G^ mutant swims as well in soft agar as WT and has neither an unwrapping nor an autoagglutination defect (not shown). Furthermore, a *flgP*^C17G^ mutant colonizes not only the chicken cecum as well as WT, but the entire chicken gastrointestinal tract. And yet, a lipidated N-terminal cysteine in FlgP is conserved across the *Campylobacterota*, with only a handful of species lacking this feature (Supp. Fig. 5).

Flagellar wrapping is a common style of swimming for polar flagellates^6^. In addition to *Campylobacter jejuni,* filament wrapping has been observed in *Shewanella putrefaciens*^48^, *Helicobacter suis*^3^, *Burkholderia* spp., and *Vibrio fischeri*^49^. Those whose flagellar motors have been imaged all have periplasmic disks. Our findings suggest a common role for the diverse and convergently-evolved periplasmic disks seen in polar flagellates

Our study demonstrates the complexity of microbial evolution that parallels that of higher eukaryotes: co-dependencies, contingencies, and enabling mutations combine to produce an interlinked system of interdependent adaptations, none of which can function in isolation.

## Acknowledgements

We thank Paul Simpson in the Imperial College London Electron Microscopy Centre for electron microscopy assistance. This work was supported by Medical Research Council grant MR/V000799/1 to EJC and MB, NIH grant R01AI065539 to DRH, and KAKENHI grant 22H05066 from the Japan Society for the Promotion of Science to DN. The funders had no role in study design, data collection and analysis, decision to publish, or preparation of the manuscript. J-JF was supported by grant TED2021-132020B-I00 from Spanish MCIN/AEI and NextGenerationEU/PRTR.

For the purpose of open access, the author has applied a Creative Commons Attribution (CC BY) license to any Author Accepted Manuscript version arising.

## Author contributions

EJC: Conceptualization, experimental design and execution, strain construction, data collection and analysis, preparation of manuscript
TD: Experimental design and execution, strain construction, data collection and analysis
DAR: Chick colonization assays
AY: Microfluidics microscopy
TU: Electron microscopy data collection
XG: Strain construction
JJF: Data analysis
EB: Holographic microscopy data collection
LW: Holographic microscopy data analysis
DN: Fluorescence microscopy data collection and analysis
DRH: Strain construction and chick colonization assays
MB: Conceptualization, experimental design, preparation of manuscript

## Materials and Methods

### Cultivation of *C. jejuni*

All strains of *C. jejuni* used in this study are derivatives of DRH212 (*rpsL*^K88R^), a streptomycin-resistant isolate of *C. jejuni* strain 81-176^50^. For all experiments, cultures were grown at 37°C on 1.4% Mueller-Hinton agar supplemented with 10 µg/mL Trimethoprim (MHT agar) plus other antibiotics as needed. Soft agar for motility assays used 0.35% MHT agar. Antibiotics were added as needed at the following concentrations: kanamycin, 50 µg/mL; chloramphenicol, 12.5 µg/mL; Streptomycin, 200 µg/mL and 2 mg/mL; Anhydrotetracycline HCl, 0.0125-0.2 µg/mL.

Solid agar, as opposed to liquid cultures, was used throughout this study in order to standardize experiments performed between the laboratories at Imperial College London, University of York and the University of Electro-Communications, as laboratories at the latter two institutions did not have access to a tri-gas incubator equipped with an orbital shaker that could be used with Class II organisms. Gas-generating sachets (Thermo-Fisher Campygen sachets (University of York) or Mitsubishi Gas sachets (UEC)) were used at these institutions.

Cell suspensions for optical microscopy experiments, tomography, and autoagglutination assays were prepared by seeding a small amount of culture from a -80°C master stock on a fresh MHT plate. Following 20-24 hours incubation, overnight growth of the inoculum was spread on another fresh MHT plate and incubated overnight. In the morning, fresh growth was gently washed off the plate into MH broth by pipetting.

### Genetic manipulation of *C. jejuni*

All mutations generated in this study are chromosomally integrated at their native loci, unless otherwise stated Strain construction was performed as previously described. Briefly, an *aphA*-*rpsL*^WT^ cassette with 500-1000 bp of flanking homology to the targeted gene was introduced by natural transformation, selecting for kanamycin resistance (Km^R^) and screening for streptomycin sensitivity (Sm^S^) on 200 µg/mL streptomycin (the *rpsL*^WT^ allele is dominant to *rpsL*^K88R^ in the merodiploid). Counterselection for loss of the *aphA*-*rpsL*^WT^ cassette was accomplished by transformation of the Km^R^ Sm^S^ intermediate strain with a fragment of DNA encoding the desired mutation, selecting for Sm^R^ on 2 mg/mL streptomycin and screening for Km^S^. Sm^R^ Km^S^ transformants were single colony purified, sequenced and stored at -80°C.

For all transformations, linear DNA was generated by SOE PCR using the protocol in Supp. file XXX, which typically yields 5-10 µg DNA/reaction. ecoRI sites were added to both ends of each fragment for methylation in order to increase transformation efficiency^51^. Typically 1-2 µg of methylated DNA was transformed for the initial Km^R^ selection, and 5-10 µg for the Sm^S^ counterselection. Excess methylated DNA can be frozen indefinitely for future strain construction projects.

### Western blotting

Western blotting was performed using whole-cell lysates. Cells were washed off plates into MH broth and the O.D._600_ adjusted to 0.5 prior to boiling in 2x SDS Laemmli buffer. 15 uL of each sample was run on 4-20% Novex wedgewell tris-glycine polyacrylamide gels. Separated proteins were transferred to 0.2 μm nitrocellulose membrane using an iBlot 2 transfer apparatus. Secondary antibodies were HRP-conjugated and imaging was performed using Clarity Western ECL substrate (Bio-rad) and a Chemidoc imaging system (Bio-rad)

Western blotting for FlgP was performed using anti-FlgP antisera raised in rabbit. Western blotting for the 6xHis-tag was performed using either HRP-conjugated primary antibody (Sigma, rabbit) or non-HRP-conjugated antisera (rabbit).

### Autoagglutination assays

For autoagglutination assays, fresh growth was washed from plates to an O.D._600_ of 1.0 in 90:10 PBS:MH broth. 1 mL of cell suspensions were pipetted into disposable polystyrene cuvettes and left to sit at ambient temperature (21-23°C) for 24 hours. O.D._600_ measurements were taken every hour for 6-8 hours, as well as a final reading at 24 hours.

In our experience, consistency is key for reproducibility in autoagglutination assays. Differences in temperature, buffer composition, agar percentage and the amount of time plates have been allowed to dry prior to use can all affect sedimentation rate. For each experiment, plates from the same batch of media were used for all strains to be compared in a given experiment. Pipetting when washing cells off overnight plates was kept to a minimum to avoid shearing flagella.

### 3D-holographic microscopy

Holographic cell tracking was performed on an inverted microscope as previously described^52^. In brief, sample chambers measuring 20 11 5 11 0.3 mm^3^ were constructed from glass slides and coverslips. These chambers were loaded with cell suspensions diluted to a concentration of approximately 3 11 10^6^ cells/ml. The standard condenser assembly in the microscope was replaced with a holder for a single-mode optical fibre directed along the optical axis of the microscope. A fibre coupled laser with a wavelength of λ=642 nm and an optical power at the sample of 3 mW/cm^2^ was used to illuminate the sample. The sample was imaged using a 2011 magnification objective lens onto a camera with pixel size of 14 um, leading to a spatial sampling frequency of 1.422 pixels/μm. Images were acquired at 100 Hz with a 3 ms exposure time. Background correction was performed by creating an image from the median pixel value at each (x,y) location, then dividing the pixel value in each frame by its corresponding value in the median image. We used Rayleigh-Sommerfeld back- propagation to create a stack of refocused images from each frame of the raw video, and segmented the corresponding 2D image stack by finding places in which the axial intensity gradient lay above a certain (manually-determined) threshold. These locations are candidates for cell positions. We then linked the coordinates in subsequent frames into cell tracks^53^, which were subjected to further analysis. Tracks shorter than 0.4 seconds were discarded. These were typically the result of cells entering and leaving the field of view. We calculated the mean-squared displacement (MSD,) for each cell^54^, and fitted the first second of data with the function . The exponent indicates the nature of the cell’s motion, and takes values between 1 (diffusive motion) and 2 (purely straight-line swimming). These values are plotted against the cells’ root-mean-squared displacement after 1 second (obtained by extrapolation for short tracks) in Figure 2.

### High-speed fluorescence microscopy

High-speed videos were recorded as described previously ^5^. Briefly, specimen chambers were prepared by adhering a 24 mm x 40 mm coverslip to a 18 mm x 18 mm coverslip using porous double-sided tape (Nichiban (size 02)). Following pipetting of sample into specimen chambers, chambers were sealed with clear nail lacquer to reduce drift.

We used DyLight 488-conjugated maleimide dye (Thermo-Fisher) to label flagellar filaments. Cell bodies were labelled using FM 4-64 dye (Life Technologies). Unless otherwise stated, all movies were captured at 400 frames per second, and cell suspensions were MH broth supplemented with methylcellulose (4000 cP, Sigma Aldrich) to a final concentration of 0.5%.

Movies were captured with an inverted microscope (IX83, Olympus), equipped with an objective lens (UPLXAPO100×OPH, N.A. 1.45, Olympus), dichroic mirrors (Di01-R488, Semrock), dual-view imaging system with optical filters (FF560-FDi01, FF03-535/50 and BLP01-568R, Semrock), a CMOS camera (Zyla 4.2, Andor), and an optical table (ASD- 1510T, JVI). Projection of the image to the camera was made at 0.065 μm per pixel. The focal position of the sample was kept at the focal position by a Z-drift compensation module (ZDC, Olympus). A blue laser beam (OBIS488-20, Coherent) was introduced into the microscope, and the resultant fluorescent images were acquired by imaging software (Solis, Andor) as 16-bit images under 2.5-ms intervals.

### Aerotaxis assays

Aerotaxis assays were performed as described previously^5^. Briefly, specimen chambers were prepared by adhering a 24 mm x 40 mm coverslip to a 18 mm x 18 mm coverslip using porous double-sided tape (Nichiban (size 02)).

Cell suspensions were adjusted to an O.D._600_ of ∼1 and pipetted into a sample chamber. Due to the speed at which populations of WT cells will aerotax, sample chambers were not sealed with clear nail lacquer. Recording was started prior to the addition of samples for the same reason.

Movies were recorded at 3 frames per second using a darkfield microscope (IX83, Olympus) equipped with an objective lens (CPLFLN10×PH, N.A. 0.3, Olympus), darkfield condenser (U-DCD, Olympus), and a CMOS camera (Zyla 4.2, Andor) and an optical table (ASD- 1510T, JVI). Projection of the image to the camera was made at 0.65 μm per pixel. Sequential images of cells were acquired by the imaging software (Solis; Andor) as 16-bit images with the CMOS camera.

Kymographs were generated in ImageJ version 1.48. The height of the sequential images was resized to one pixel and aligned vertically so that the y-axis represents time.

### Microfluidic experiments

Microfluidic devices with confined 1 μm channels were fabricated using standard photolithography and soft lithography methods as described previously^55^. Briefly, polydimethylsiloxane (PDMS, Sylgard 184, Dow), a two-part silicone elastomer, was cast over a photolithography master and cured at room temperature for 48 h. A piece of PDMS was cut out using a scalpel and used as a microfluidic device. Cell suspensions with MH broth containing 0.5% methylcellulose, were dropped onto a glass slide and then covered with the microfluidic device casting from the top. Movies were captured with an inverted microscope (IX73, Olympus), equipped with an objective lens (UPLXAPO100×OPH, N.A. 1.45, Olympus), a filter set (GFP-4050B, Semrock), mercury lamp (U-HGLGPS, Olympus), a CMOS camera (DMK33UX174, Imaging Source), and an optical table (HAX-0806, JVI). Projection of the image to the camera was made at 0.058 μm per pixel. Sequential images were acquired by the imaging software (Solis, Andor) as 16-bit images under 25-ms intervals.

### Electron cryotomography and subtomogram averaging

Strains to be imaged for subtomogram averaging were washed off plates and concentrated to an O.D._600_ of 10-20 and mixed with 10 nm gold fiducial markers (Sigma-Aldrich) in 5% BSA. Samples were applied to Quantifoil R2/2 grids and plunge frozen in liquid ethane using a Vitrobot (FEI). Imaging was performed on a Thermo-Fisher Glacios 200 kV electron microscope equipped with a Falcon 4 direct electron detector and Selectris energy filter.

Tomograms were reconstructed using a combination of IMOD 4.11.8 for fiducial modeling and Tomo3D for SIRT tomographic reconstruction^56,57^. To enhance the contrast of tomograms for display of unaveraged motors and to measure disk diameters, tomograms were CTF-deconvoluted as first described by (Tegunov and Cramer 2019)^58^ but with CTF deconvolution performed in 2-D on the tilt series prior to 3-D tomographic reconstruction. In short, the procedure restores the magnitude of the low-resolution components that are attenuated by the CTF while removing the noisy components at medium and high resolution, which results in an overall contrast improvement. This code is available in version 2.2 of Tomo3D.

For subtomogram averaging, particles were picked using 3dmod from the IMOD suit and imported into Dynamo 1.1.532 for subtomogram averaging. We imposed C17 symmetry for averaging based upon established prominent symmetry of the periplasmic structures of the *C. jejuni* motor. STA maps have been deposited in the EMDB with the following accession codes: EJC168 12.5 ng/mL ATc - EMD-XXXXX; EJC168 25 ng/mL ATc - EMD-XXXXX; EJC168 50 ng/mL ATc - EMD-XXXXX; EJC168 100 ng/mL ATc - EMD-XXXXX; *flgP*^AAA^ - EMD-XXXXX; *flgP*-*lpp*^55^ - EMD-XXXXX; Δ*flgPQ* - EMD-XXXXX; EJC103 - EMD-XXXXX; *flgP*^Δ18–62^ - EMD-XXXXX.

### Chicken colonization assays

Chick colonization assays. The ability of WT C. jejuni 81-176 rpsLSm and isogenic mutants to colonize chicks after oral inoculation was determined as previously described (32). Briefly, fertilized chicken eggs (SPAFAS) were incubated for 21 days at 37.5 °C with appropriate humidity and rotation in a Digital Sportsman model 1502 incubator (Georgia Quail Farms Manufacturing Company). One day after hatch, chicks were orally inoculated with 100 μL of phosphate buffered saline (PBS) containing approximately 180-240 CFU WT or mutant strains. Strains were prepared for infection after 16 h growth at 37 °C under microaerobic conditions on MH agar by suspending C. jejuni strains in MH broth. Dilution series in PBS were performed to achieve the appropriate inoculum for oral gavage of chicks. Dilutions of the inoculum were plated on MH agar to assess the number of bacteria in each inoculum. At 7 days post-infection, chicks were sacrificed, the cecal contents were removed and suspended in PBS, and serial dilutions were plated on MH agar containing trimethoprim and cefoperazone. Following 72 h of growth at 37 °C in microaerobic conditions, bacteria were counted to determine CFU per gram of organ content. Recovered colonies were analyzed by colony PCR to verify that WT and mutant strains were isolated from respectively infected chicks.

### Whole genome sequencing

Whole genome sequencing was performed by Source Biosciences (U.K.). Genomes were assembled and analysed using the software package Geneious Prime 2021.0.3 (Biomatters, New Zealand). The paired reads provided by Source Biosciences were imported into Geneious and trimmed using BBDuk, removing adapters and low-quality reads. Whole genome sequencing reads of parental strains *flgP-lpp*^56^ and Δ*flgPQ* were mapped to a *C. jejuni* reference genome NC_008787. These assembled genomes were then used as reference genomes against which suppressor genomes were assembled and analysed. We used the Geneious variant finder to find mutations in each sequenced suppressor genome relative to its parental reference genome, characterise mutation frequency and its possible effect on codon and amino acid changes.

### Phylogenetics

A phylogenetic tree of the *Cjj_81176_0661* family was determined using the sequences allocated to the PFAM PF04748 family as downloaded on 17th May 2022 and performing a multiple sequence alignment using RAxML with a Jones-Taylor-Thornton (JTT) model of amino acid substitution rates with a discrete gamma distribution.

---

**Supplemental Figure 1:**
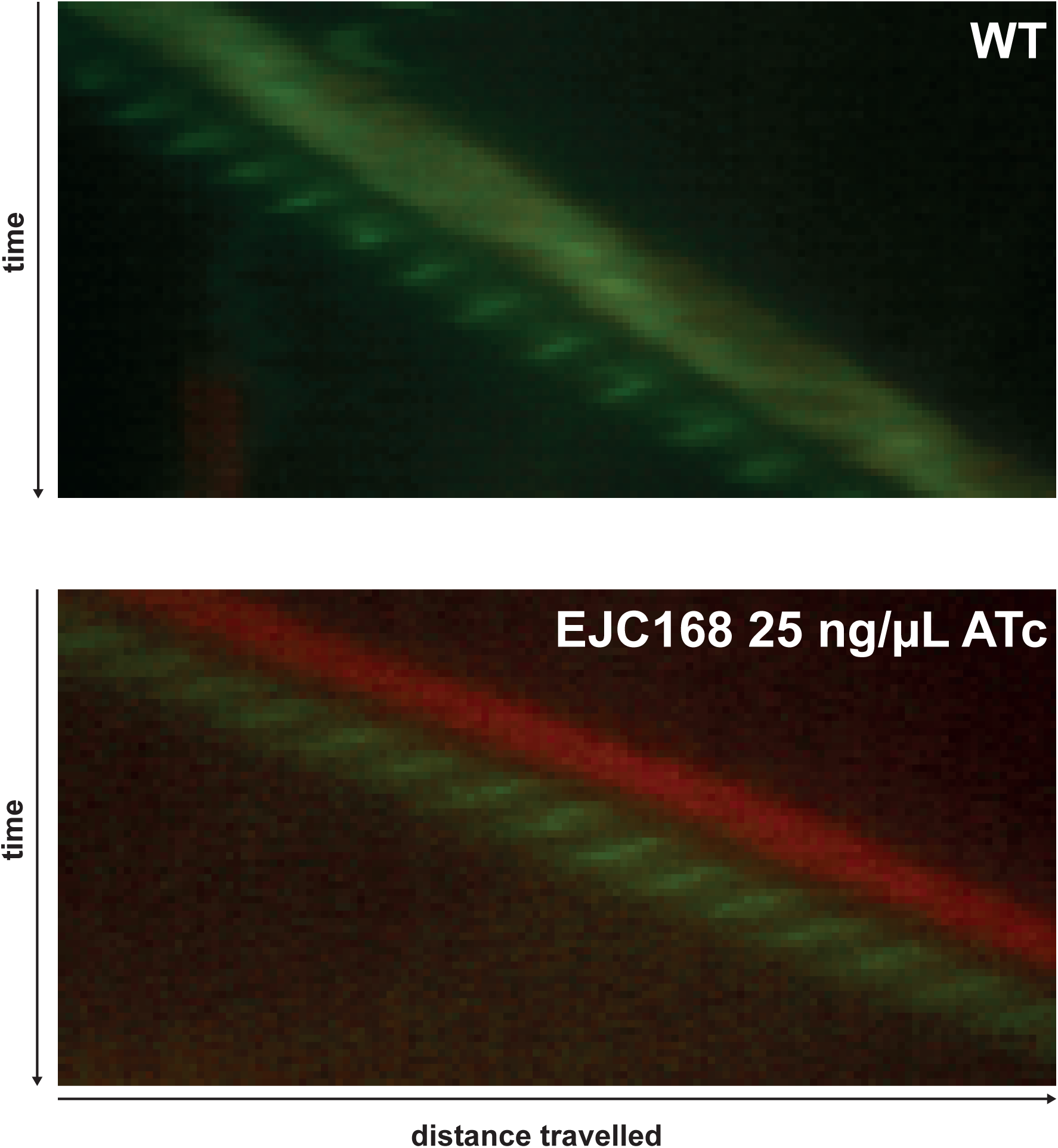
Smaller basal disks do not impair motor rotation. Comparison of motor-rotation rate of WT cells and EJC168 under low induction conditions demonstrated that motors small-diameter basal disks rotated at rates comparable to WT motors.

**Supplemental Figure 2:**
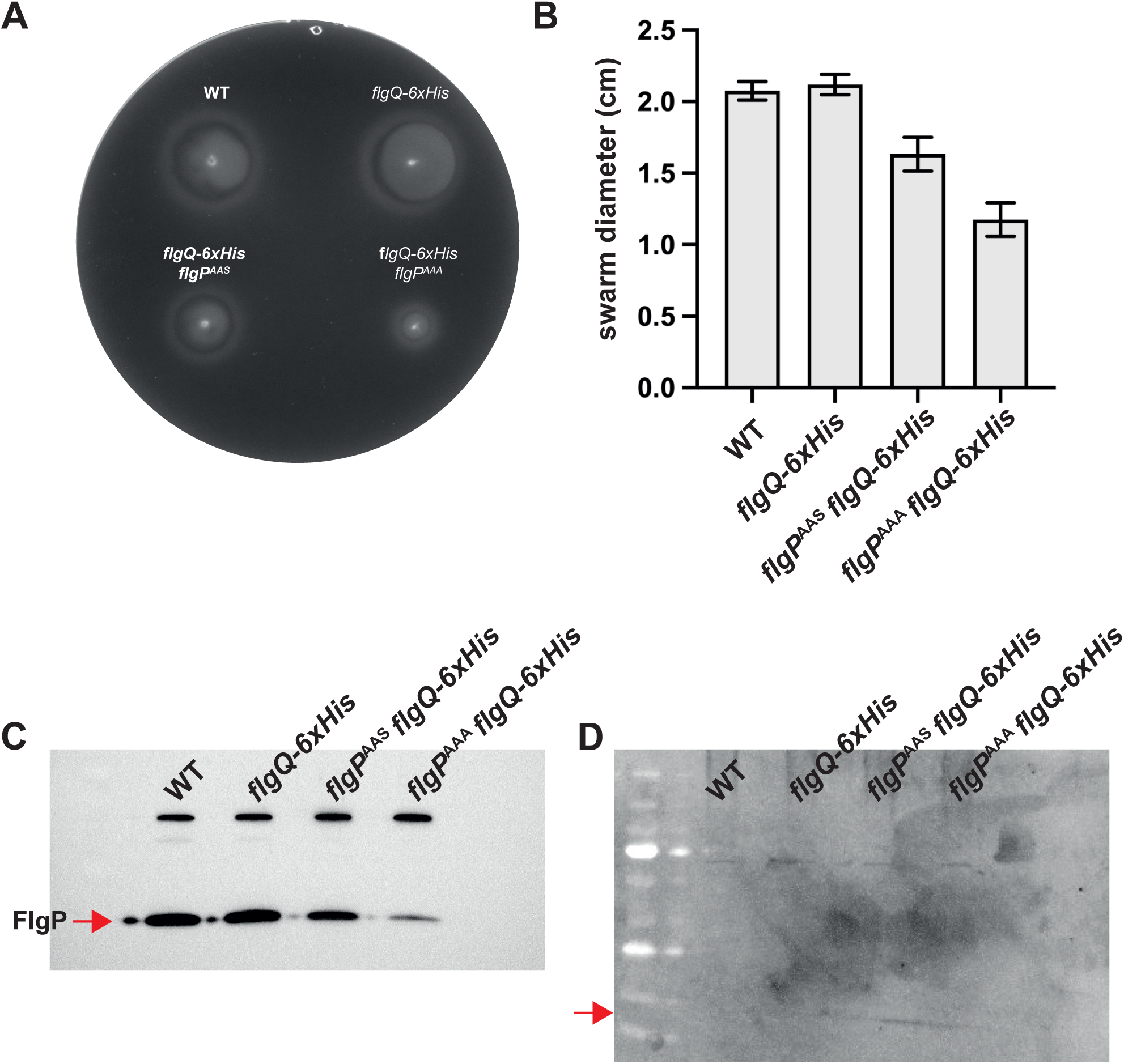
The *flgP*^AAA^ mutant expresses lower levels of FlgP. The *flgP*^S69A^ ^E157A^ ^K159S^ (*flgP*^AAS^) and *flgP*^S69A^ ^E157A^ ^K159A^ (*flgP*^AAA^) mutants were constructed in a *flgQ-6xHis* background **(A** and **B)** in order to determine whether any observed phenotypes arising from the alanine-scanning mutations in *flgP* were due to polarity effects on downstream *flgQ*. **(C)** Western blotting for FlgP revealed that lower levels of FlgP are produced in both *flgP*^AAS^ and *flgP*^AAA^. That both mutants have amino-acid substitutions at identical sites, but exhibit different levels of FlgP by western blot that correlate with their different motility levels in soft agar indicates that the reduced intensity of the FlgP band in *flgP*^AAA^ is not due to reduced epitope binding by the anti-FlgP antisera. **(D)** Multiple attempts at western blotting for FlgQ-6xHis were unsuccessful, as no expected bands were observed (arrow) for any strain tested. Thus, we can not rule out that the *flgP*^AAS^ and *flgP*^AAA^ alleles impact *flgQ* expression at this time.

**Supplemental Figure 3:**
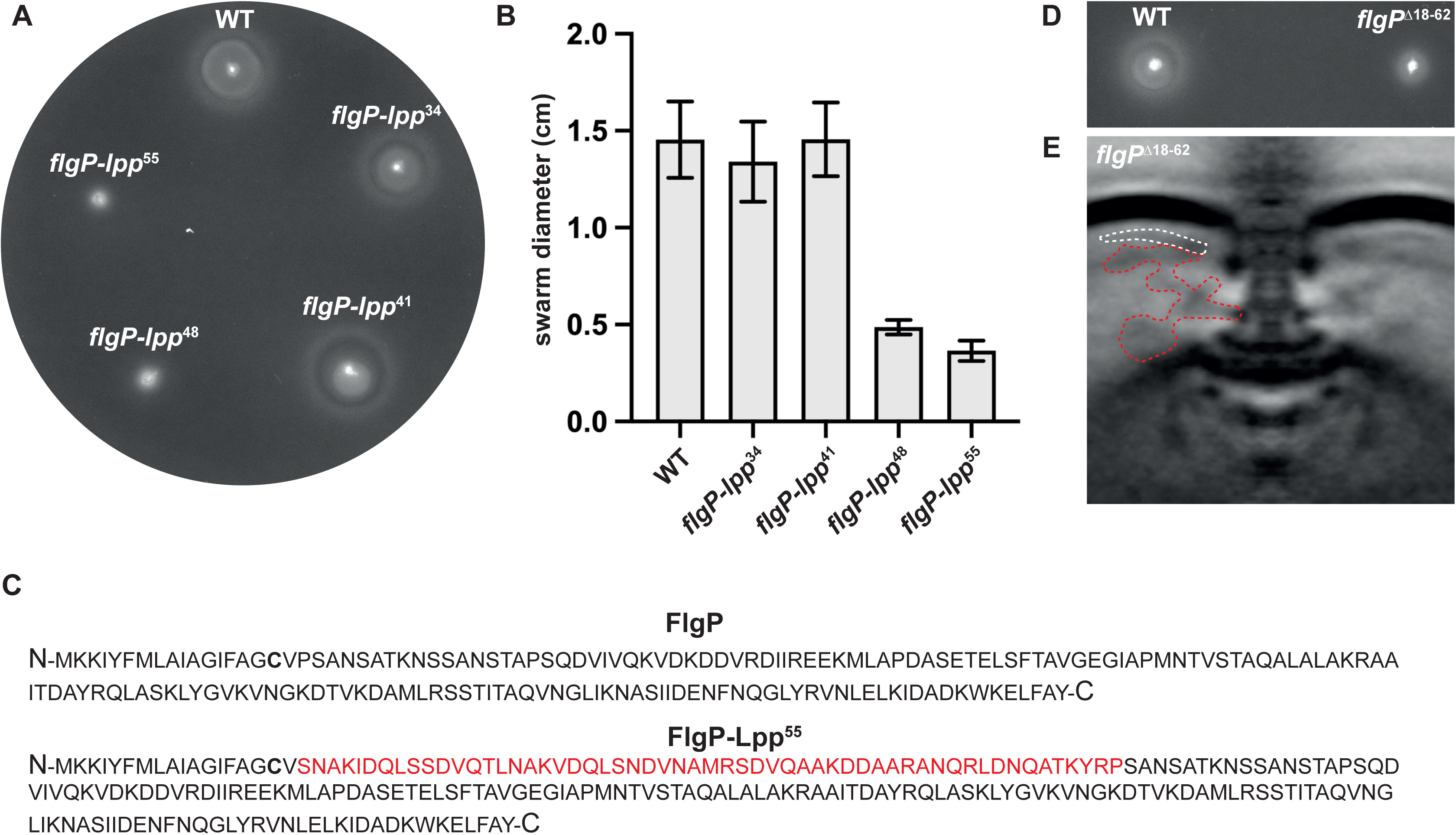
Lengthening or shortening the FlgP linker displaces the basal disk and impacts motility. **(A** and **B)** Lpp-linkers of increasing length were inserted at the N-terminus of the FlgP OM- linker. Linkers ≥ 48 residues resulted in a significant decrease in soft-agar swarm diameter. **(C)** The *flgP-lpp* mutant that exhibited the poorest motility in swarm diameter, *flgP-lpp*^55^, was selected for further investigation. **(D)** Deletion of the FlgP OM-linker, leaving C17 in place (*flgP*^Δ18–62^), decreased but did not abolish motility. **(E)** The basal disk (white dashed line) is pulled toward the OM, and out of register with the P-ring, in the *flgP*^Δ18–62^ mutant. The stator scaffolding (red dashed line) is also distorted and is appears to be unstably associated with the motor in this background.

**Supplemental Figure 4:**
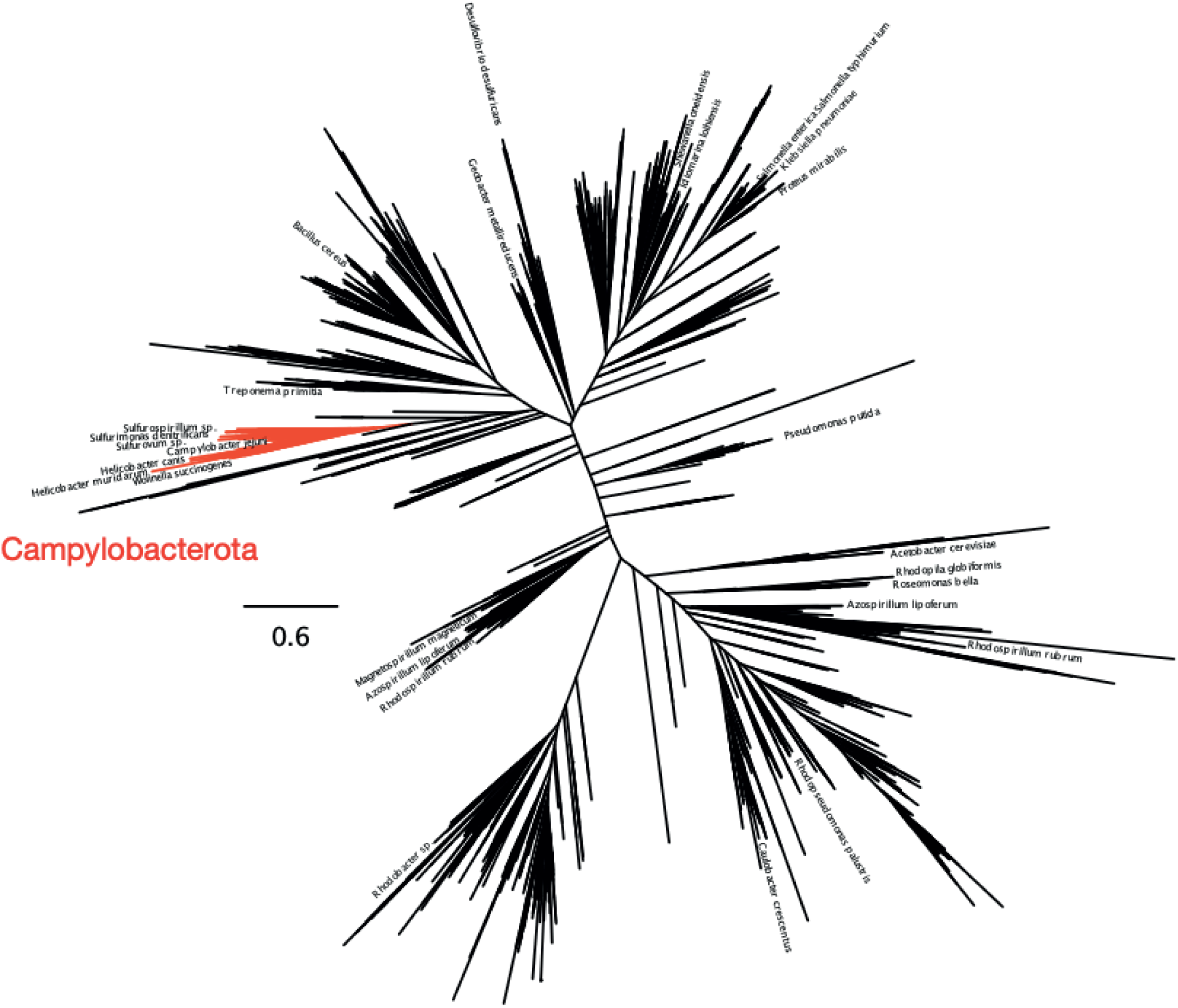
*Cjj_81-176_0661* is a predicted polysaccharide deacetylase found only in the Campylobacterota. *Cjj_81-176_0661* (*661*) encodes a protein that is a member of the PF04748 divergent polysaccharide deacetylase family. *661* is ubiquitous in the Campylobacterota but is not found in other genera. The ubiquity of *661* in the Campylobacterota suggests an ancient origin at the base of the Campylobacterota tree.

**Supplemental Figure 5:**
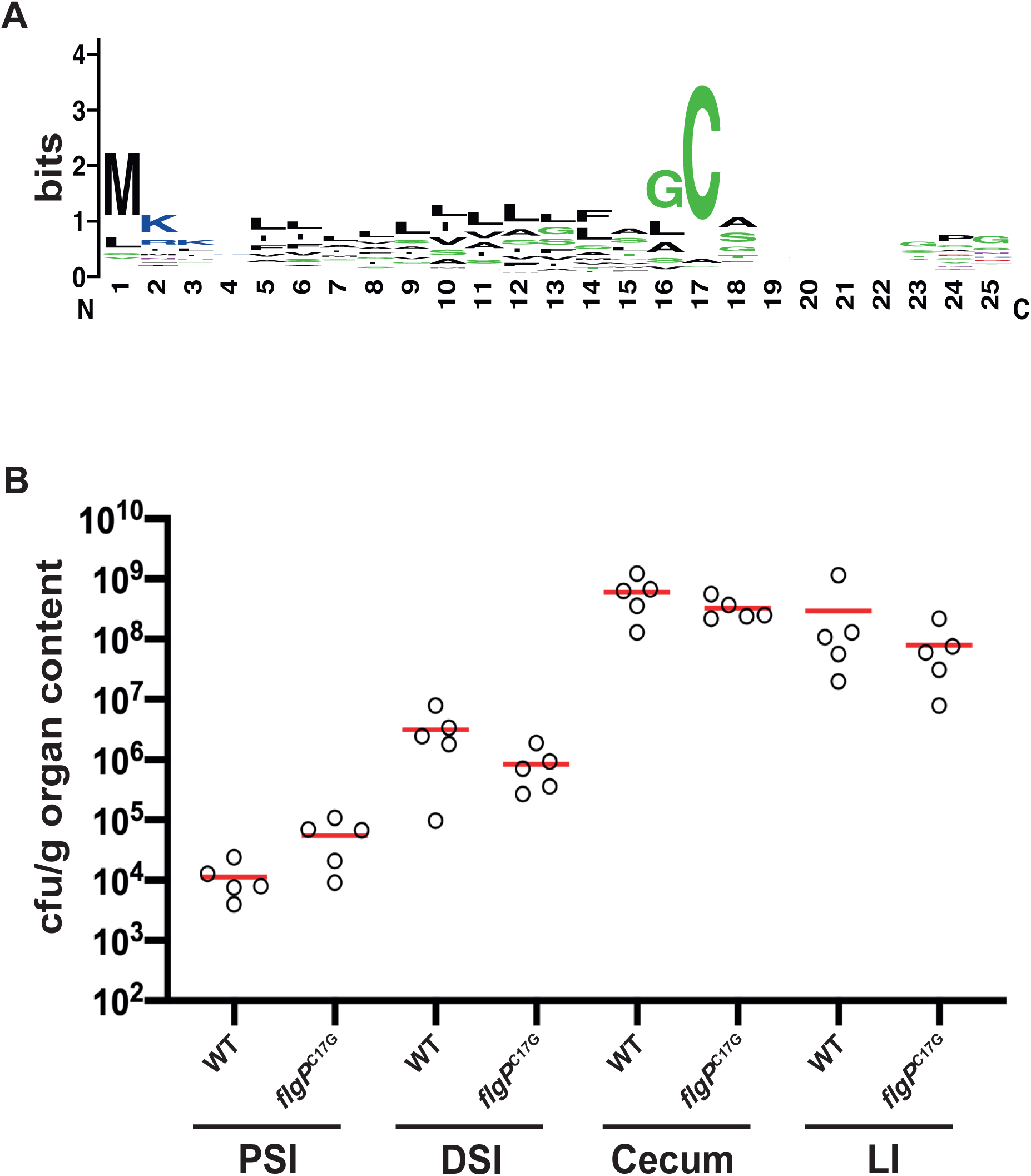
An N-terminal cysteine is unimportant for host colonisation, but is a conserved feature across the Campylobacterota genus. **(A**) The N-terminal residues of FlgPs from 71 members of the Campylobacterota were aligned, revealing that an N-terminal, presumably lipidated cysteine is a conserved feature of the protein. **(B)** Comparison of colonisation ability of WT and a *flgP*^C17G^ mutant along the chick gastrointestinal tract did not reveal a colonisation defect in the *flgP*^C17G^ background (PSI: proximal small intestine; DSI: distal small intestine; LI: large intestine).

**Supplemental video 1:** 10x magnification darkfield video of WT cells in a sample chamber.

**Supplemental video 2:** 10x magnification darkfield video of EJC168 under low (25 ng/mL ATc) induction of *flgPQ*.

**Supplemental video 3:** Video of autoagglutinated clump of low-induction EJC168 showing cells in the clump are motile but unable to overcome filament-filament binding.

**Supplemental video 4:** Video of a *flgP*::*lpp*^55^ cell swimming. The failure of the cell to unwrap during motor reversals results in a stuttering type of motility. This video is the source of the kymograph in figure 2E (bottom panel).

**Supplemental video 5:** Video of WT cell swimming. Upon directional reversal of motor rotation, the cell swaps which filament is wrapped, leading to reversal of swimming direction. This video is the source of the kymograph in figure 2E (top panel).

**Supplemental video 6:** 10x magnification darkfield videos (false colored) comparing aerotactic behaviour of a populations of WT cells to *flgP*::*lpp*^55^ cells.

**Supplemental video 7:** Video of cells entering and exiting a PDMS microfluidic device with 1 µm channels. Filaments were labelled with DyLight 488-conjugated maleimide dye.

**Supplemental videos 8 and 9:** Videos of *flgP*::*lpp*^55^ cells swimming through a channel of a PDMS microfluidic device. Due to their inability to unwrap their filaments, pile-ups of cells occur in the channels in the presence of an obstacle. Videos 8 and 9 are sequential, observing the same pile-up over the course of a couple minutes.

**Supplemental video 10:** Video of WT cells swimming through a channel of a PDMS microfluidic device. WT cells are able to reverse direction upon encountering an obstacle due to their ability to unwrap their filaments from the cell body.

**Supplemental video 11:** Triptych video comparing Δ*flgPQ*, Δ*flgPQ* Δ661 and Δ*flgPQ* Δ*kpsD* Δ*pglAB*::*aphA* cells. Deletion of *661* results in more cells with rotating filaments compared to the Δ*flgPQ* single mutant.

